# Apoptotic caspases cleave DRP1 to promote mitochondrial fusion and anti-viral immune responses

**DOI:** 10.1101/2024.07.13.603409

**Authors:** Yujie Fang, Zihan Guan, Xiangtao Zhu, Zhenqiong Guan, Shufen Li, Ke Peng

**Affiliations:** Basic Medical College, Changzhi Medical College, Changzhi, Shanxi, 046000, China; State Key Laboratory of Virology and Biosafety, Center for Antiviral Research, Center for Biosafety Mega-Science, Wuhan Institute of Virology, Chinese Academy of Sciences, Wuhan 430207, China; University of Chinese Academy of Sciences, Beijing 100049, China; Provincial Key Laboratory of Jiangxia, Wuhan, 430207, Hubei

**Keywords:** Caspase, DRP1, mitochondrial elongation, innate immunity, virus

## Abstract

Apoptosis has been recognized as a suicidal host-defense programmed cell death pathway against invading pathogens. However, recent evidences showed that viruses can employ caspases to cleave and inactivate immune signaling molecules to facilitate infection. Whether caspases can instead promote anti-viral immune responses is currently unknown. Here, we demonstrated that the nonstructural protein (NSs) of Rift Valley fever virus (RVFV) triggers activation of apoptotic caspases, which cleave the mitochondrial fission factor dynamin-related protein 1 (DRP1) resulting in mitochondrial elongation. Elongated mitochondria promote mitochondrial antiviral-signaling protein (MAVS) aggregation leading to enhanced anti-viral immunity. Apoptotic caspases, including caspase-3, -6, -7 and -8, cleave DRP1 at the motifs of D^500^FAD^503^ and/or AEAD^556^, suggesting that this cleavage event may occur during infection of different viruses. Indeed, infection of influenza A virus subtype H1N1, Sendai virus (SeV) and herpes simplex virus 1 (HSV-1) all triggered apoptotic caspases activation to cleave DRP1 promoting anti-viral immune responses. Compared with wild-type DRP1, introduction of caspase-resistant DRP1 strongly attenuated immune responses triggered by virus infection. These results revealed a mechanism through which apoptotic caspases promote anti-viral immunity by regulating mitochondrial morphodynamics.

## Introduction

Rift Valley fever virus (RVFV) is an enveloped, single-stranded ambisense, tri-segmented RNA virus, belonging to the genus *Phlebovirus* and the family *Phenuiviridae* within the order *Bunyavirales* (Noronha and Wilson, 2017). As a mosquito-borne virus, RVFV is transmitted among humans through mosquito bites and can cause serious diseases including ocular (eye) disease, meningoencephalitis or haemorrhagic fever (McMillen and Hartman, 2018). The case-fatality ratio for RVF patients developing the haemorrhagic fever can be up to approximately 50% (Sun et al., 2018). Due to the absence of effective preventive or therapeutic strategies for human use and the potential of widespread transmission, RVFV has been listed as one of the viruses causing priority diseases categorized by the World Health Organization (WHO) (Gerken et al., 2022). RVFV has a three-segmented RNA genome of the L (large), M (medium) and S (small) segment, of which the S segment encodes the major virulence factor nonstructural protein (NSs) (Freire et al., 2015; Leger et al., 2020). Previous studies have shown that the virulence factor NSs protein triggers apoptosis during infection and contributes to viral pathogenesis (Austin et al., 2012; Narayanan et al., 2014; Smith et al., 2010).

Apoptosis is a form of programmed cell death that functions in regulating normal embryonic development, tissue homeostasis and immune responses (Elmore, 2007; Everett and McFadden, 1999; Williams, 1994). Apoptosis is mediated through the activation of apoptotic caspases including caspase-2, -3, -6, -7, -8, -9 and -10. Apoptosis can be triggered by either extrinsic or intrinsic pathways. The extrinsic apoptosis pathway is activated by the engagement of cell surface death receptors, such as members of the tumor necrosis factor (TNF) receptor superfamily, by its ligands, leading to the activation of caspase-8 and its downstream effector caspases (Li and Yuan, 2008; Orzalli and Kagan, 2017; Zhou et al., 2017). The intrinsic apoptosis is regulated by the function of pro-apoptotic members of the B-cell lymphoma 2 (Bcl-2) family, which controls the Bax/Bak activation and induces mitochondrial outer membrane permeabilization (MOMP) (Zhou et al., 2017). MOMP results in the release of cytochrome c into the cytosol, which binds to Apaf-1 to activate caspase-9 and the downstream effector caspases-3, -6 and -7 (Li and Yuan, 2008; Zhou et al., 2017).

Many viruses, including both DNA and RNA viruses, cause infected cells to undergo apoptosis (Koyama et al., 1998). Apoptosis has long been regarded as a host defense mechanism eliminating the niche of virus amplification (Koyama et al., 1998). However, recent evidences showed that viruses can employ activated caspases to cleave and inactivate the innate immune molecules to dampen the host anti-viral immune responses and promote virus replication (Ning et al., 2019; Wang et al., 2017). For example, vesicular stomatitis virus (VSV) and Poliovirus trigger caspase-3 or caspase-7 activation to cleave and inactivate mitochondrial antiviral-signaling protein (MAVS) (also named as VISA, IPS-1, and Cardif) to facilitate virus replication (Ning et al., 2019; Rebsamen et al., 2008). Similarly, human immunodeficiency virus type 1 (HIV-1) triggers caspase-3 activation to cleave interferon regulatory factor 3 (IRF3) and promotes virus infection (Ning et al., 2019; Park et al., 2014). IRF3 can also be cleaved and inactivated by caspase-8 during Sendai virus (SeV) infection (Sears et al., 2011). Up till now, whether virus infection-triggered apoptotic caspases can promote antiviral innate immunity has not been reported.

Both extrinsic and intrinsic apoptosis involve regulation of mitochondria, a critical intracellular organelle participating in energy production, metabolism, cell death and differentiation (Ren et al., 2020). Recent studies revealed that mitochondria are central hubs of innate immune responses mediating both type I interferon (IFN-I) and inflammatory responses (Sandhir et al., 2017). Mitochondria undergo dynamic morphological changes regulated by mitochondrial fusion and fission factors (Tilokani et al., 2018). Key regulators that mediate the switch between mitochondrial fusion and fission include optic atrophy 1 (OPA1), which is responsible for fusion of the inner membrane; mitofusin 1 (MFN1) and mitofusin 2 (MFN2), which are responsible for fusion of the outer membrane; dynamin-related protein 1 (DRP1/DNM1L/DLP), which is the critical factor that participates in mitochondrial fission (Chen et al., 2020; Ren et al., 2020; Wei et al., 2021); and mitochondrial fission factor (MFF) and mitochondrial fission protein 1 (FIS1), which serve as accessory adaptors that facilitate the recruitment and function of DRP1 (Ren et al., 2020). The morphodynamics of mitochondria is important for regulating immune responses with the elongated mitochondria mediates enhanced immune responses through serving as the platform for immune signalosome assembly (Chen et al., 2020). Mitochondrial elongation has been reported during virus infection which is induced by inactivation of DRP1 modulated by its phosphorylation status (Chen et al., 2020). Whether virus-triggered apoptosis can regulate mitochondrial morphodynamics is currently unknown.

Here, we found that RVFV virulence factor NSs activates apoptotic caspases which then cleave and deplete mitochondrial fission factor DRP1. DRP1 depletion leads to elongated mitochondria and promotes anti-viral immune responses through facilitating MAVS aggregation. We further demonstrated that multiple apoptotic caspases can cleave and deplete DRP1 at different positions. This caspase mediated DRP1 cleavage event is observed during both DNA and RNA virus infection and appears as a general caspase-dependent host anti-viral mechanism.

## Results

### RVFV infection leads to mitochondrial elongation through down-regulating DRP1

RVFV infection leads to distinctly elongated mitochondria in the infected PMA-induced THP-1 macrophage cells (THP-1^PMA^) (Figure 1A). The RVFV-induced mitochondrial elongation was also observed in various cell types including Huh7 cells and A549 cells (Figure 1A). Examination of ultrathin sections of RVFV infected THP-1^PMA^ cells by transmission electron microscopy (TEM) confirmed that mitochondria were more elongated in RVFV-infected cells than those in uninfected cells (Figure 1B). To identify which viral protein induced mitochondrial elongation, RVFV viral proteins, including NSs, NP, NSm, Gn, Gc and polymerase (Pol/RdRp), were transiently expressed in Huh7 cells and the mitochondrial morphology was observed. It is found that only NSs expression, but not other viral proteins, induced distinct mitochondrial elongation (Figure 2A). The expression of RVFV-NSs in A549 cells also induced mitochondrial elongation (Figure S1A). To analyze whether NSs mediates mitochondrial elongation during RVFV infection, we generated NSs-deleted RVFV encoding EGFP in the position of NSs (RVFV^EGFP △ NSs^) and evaluated mitochondrial morphology in infected cells. Unlike infection with wild type (WT) virus, infection with RVFV^EGFP△NSs^ did not lead to elongated mitochondria (Figure 2B). These results suggested that the viral protein NSs induced mitochondrial elongation during RVFV infection.

**Figure 1.**
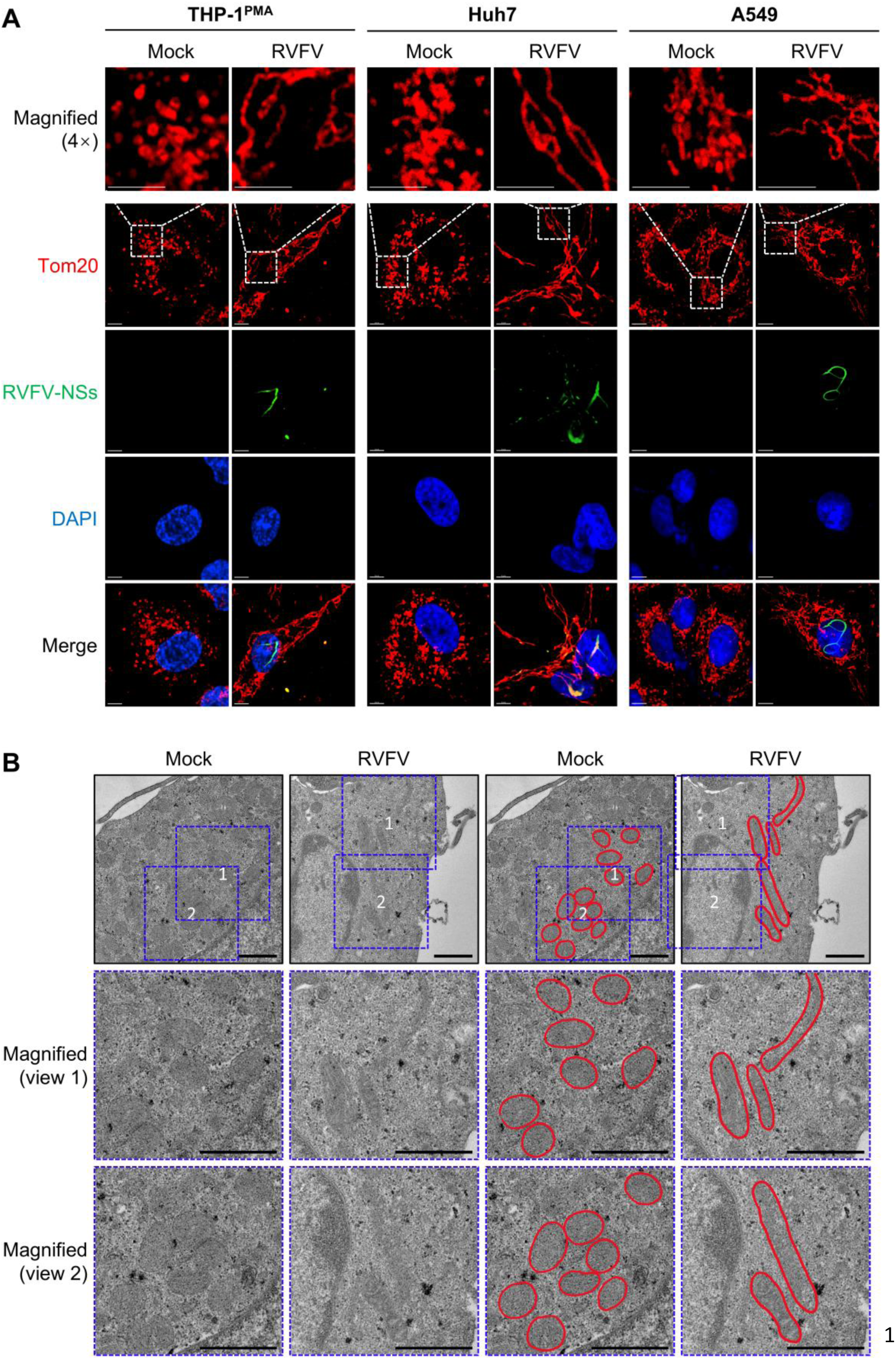
RVFV infection leads to mitochondrial elongation. (A) Mitochondria were visualized by immunofluorescence using confocal microscopy. THP-1^PMA^, Huh7 or A549 cells were infected or mock infected with RVFV for 24 h at MOI = 5. Mitochondria were labeled with anti-Tom20 antibody (red), RVFV were labeled with anti-RVFV-NSs antibody (green) and Nuclei were stained with DAPI (blue). Scale bars, 5 μm (THP-1^PMA^ and Huh7 cells) or 7.5 μm (A549 cells).(B) Sections of mock infected or RVFV-infected THP-1^PMA^ cells (MOI = 1) were analyzed by TEM. Scale bars, 1 μm. Blue dashed squares represent the magnification of regions of interest. Mitochondria are circled in red shown on the right panels. Data are representative of three experiments with similar results.

**Figure 2.**
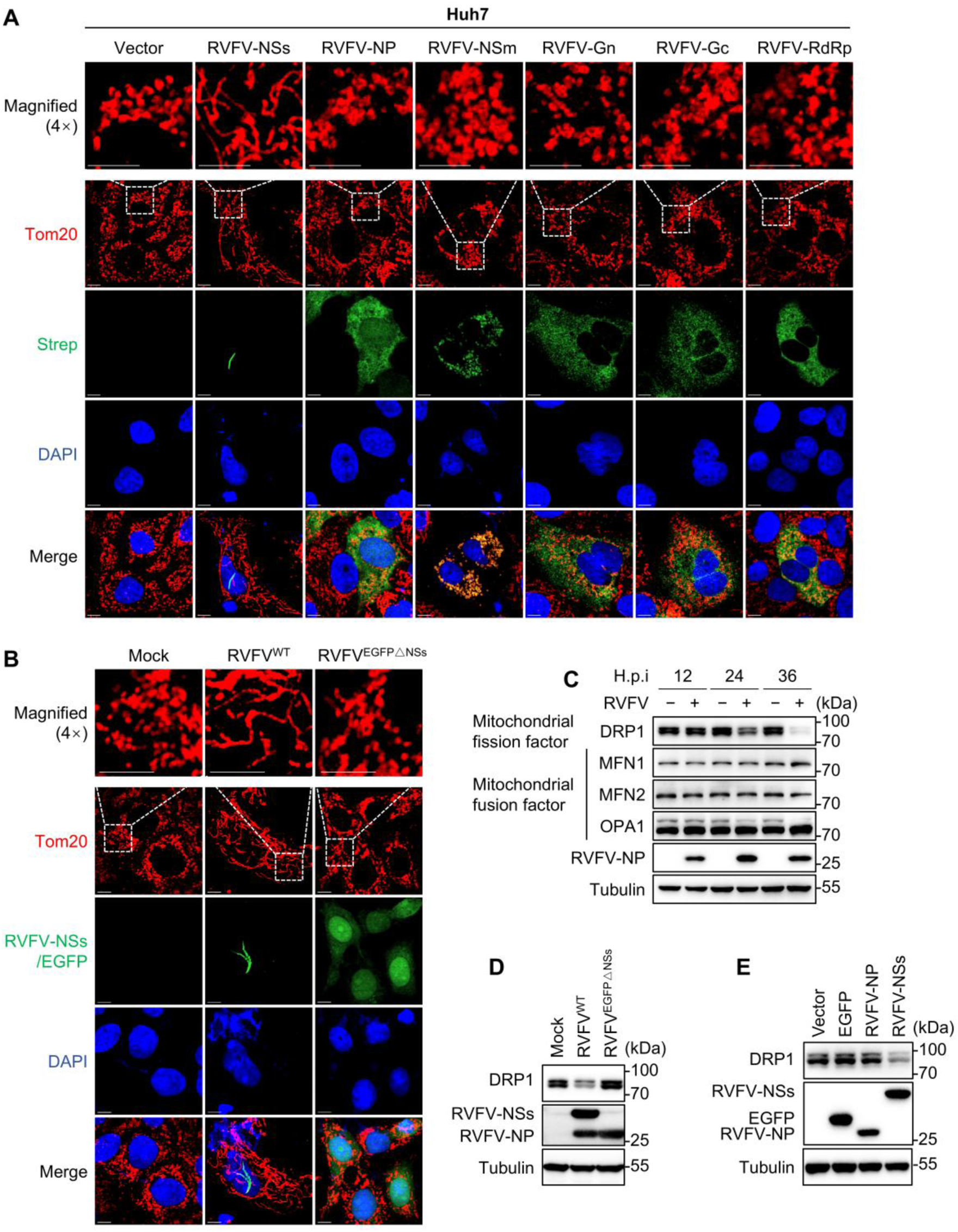
RVFV infection induces NSs-dependent DRP1 downregulation leading to mitochondrial elongation. (A) The effects of exogenously expressed RVFV viral proteins on mitochondrial morphology in Huh7 cells was observed by confocal microscopy. Huh7 cells were transfected with empty vector or vectors expressing strep-tagged viral proteins of RVFV for 48 h. Mitochondria were labeled with anti-Tom20 antibody (red), viral proteins of RVFV were labeled with anti-strep antibody (green) and Nuclei were stained with DAPI (blue). Scale bars, 7.5 μm. (B) Huh7 cells were mock infected or infected with the indicated virus for 24 h. Mitochondria were labeled with anti-Tom20 antibody (red), RVFV were labeled with anti-RVFV-NSs antibody (green), RVFV^EGFP△NSs^ were observed through detecting EGFP signal, and Nuclei were stained with DAPI (blue). Scale bars, 7.5 μm. (C) Immunoblot analysis of the proteins that regulate mitochondrial morphology, including MFN1, MFN2, OPA1 and DRP1 in THP-1^PMA^ cells infected with RVFV for the indicated times. (D) Immunoblot analysis of DRP1 protein level in Huh7 cells infected or mock infected with RVFV^WT^ or RVFV^EGFP△NSs^ for 24 h. (E) Immunoblot analysis of DRP1 protein level in Huh7 cells transfected with the plasmids expressing the indicated proteins for 48 h. Data are representative of three experiments with similar results.

To explore the mechanisms underlying RVFV-NSs induced mitochondrial elongation, we first examined the abundance of host proteins that regulate mitochondrial morphology, including MFN1, MFN2, OPA1, which mediate mitochondrial fusion, and DRP1, which mediates mitochondrial fission. It is found that among these proteins the fission factor DRP1 was significantly down-regulated in RVFV infected Huh7 cells, while the abundance of other proteins was unaffected (Figure 2C). Additionally, DRP1 down-regulation in RVFV infected cells showed a MOI-dependent phenotype (Figure S1B). DRP1 down-regulation was also observed in RVFV infected human A549 cells (Figure S1C), mouse bone marrow-derived macrophages (BMDMs) (Figure S1D) or liver samples harvested from RVFV-infected mice (Figure S1E). Since DRP1 functions in mitochondrial fission, it’s down-regulation might account for mitochondrial elongation triggered by RVFV infection. In line with this notion, infection with RVFV^EGFP △ NSs^, which does not result in mitochondrial elongation (Figure 2B), did not lead to down-regulation of DRP1 in Huh7 cells (Figure 2D). Furthermore, exogenous expression of NSs leads to down-regulation of DRP1 protein level (Figure 2E). Taken together, these results suggested that RVFV infection induced NSs-dependent DRP1 down-regulation resulting in mitochondrial elongation.

### DRP1 is cleaved by activated caspase during RVFV infection

We next investigated the molecular mechanisms underlying RVFV NSs-induced DRP1 down-regulation. The transcriptional level of DRP1 was not down-regulated during RVFV infection (Figure S2A), indicating down-regulation at the post-transcriptional level. Since cellular protein degradation can occur through the ubiquitin-proteasome system, the autophagy-lysosome pathway, and caspase-mediated cleavage, we evaluated these pathways respectively. The MITOL-mediated DRP1 ubiquitylation and degradation was reported before (Das et al., 2022), we therefore first assessed the role of the ubiquitin-proteasome system using the proteasome inhibitor MG132. To validate MG132 efficacy, we detected the levels of PKR in RVFV infected cells, a known target of RVFV-induced proteasomal degradation (Mudhasani et al., 2016). Our results showed that while MG132 treatment effectively inhibited the RVFV-induced PKR degradation, it did not inhibit RVFV-induced DRP1 downregulation (Figure 3A). We then examined the autophagy-lysosome pathway, and found that treatment with chloroquine (CQ), an autophagy-lysosome inhibitor, also did not significantly prevent DRP1 down-regulation in the RVFV infected cells (Figure 3B). In contrast, the pan caspase inhibitor Z-VAD-FMK treatment prevented RVFV-induced DRP1 down-regulation in a dose-dependent manner (Figures 3C and 3D). These results suggested that RVFV infection triggered the caspase-mediated down-regulation of DRP1.

**Figure 3.**
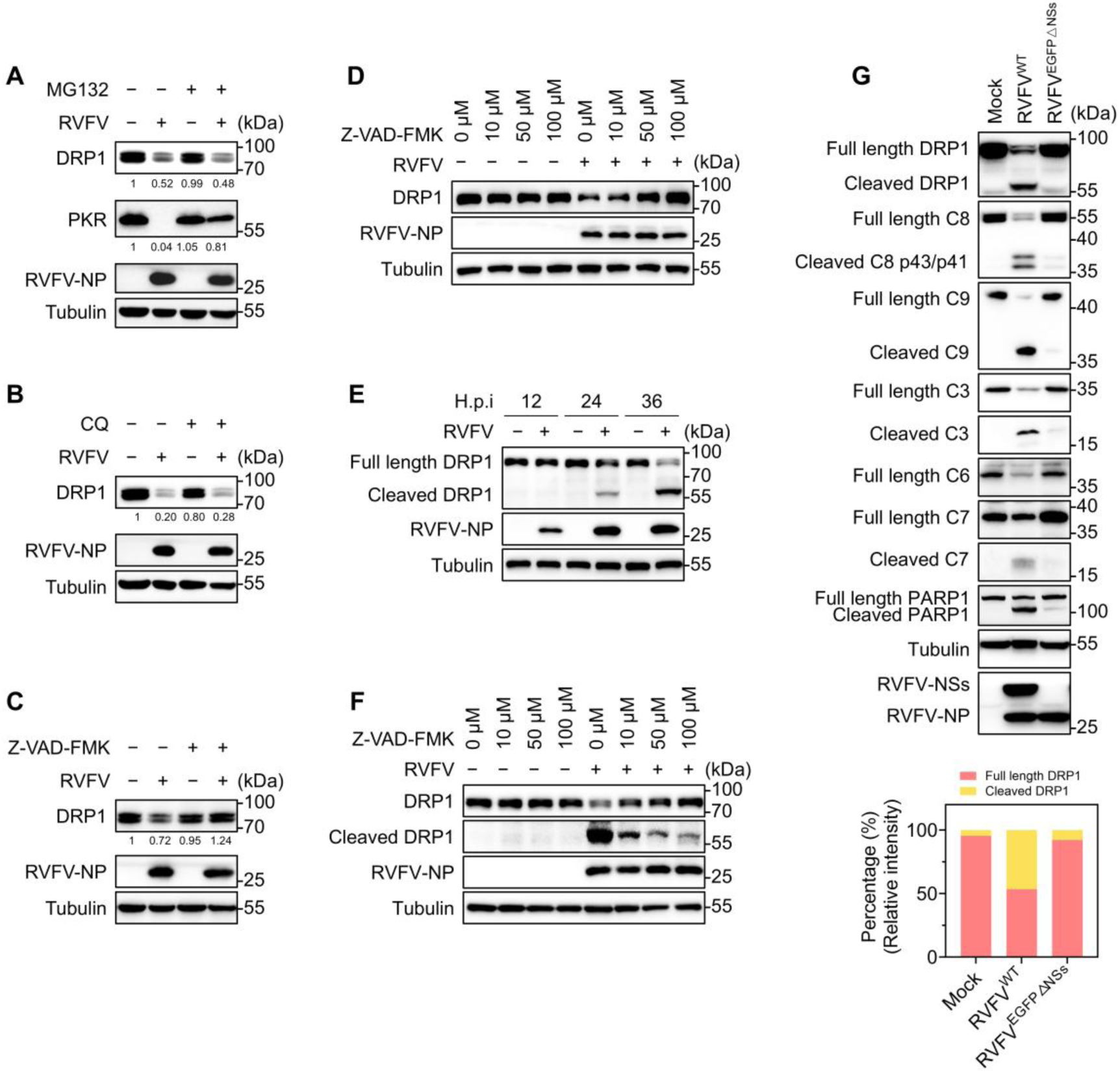
DRP1 is cleaved by activated caspase during RVFV infection. (A-C) Immunoblot analysis of indicated protein levels in Huh7 cells infected or mock infected with RVFV for 2 h, followed by MG132 (5 μM) (A), CQ (10 μM) (B) or Z-VAD-FMK (20 μM) (C) treatment for 22 h. The numbers below each lane represent protein levels relative to Tubulin and expressed relative to the control group (defined as 1) by ImageJ gray value analysis. (D) Immunoblot analysis of DRP1 protein level in THP-1^PMA^ cells infected or mock infected with RVFV for 2 h, followed by Z-VAD-FMK treatment with the indicted concentration for 22 h. (E) Immunoblot analysis of DRP1 cleavage in THP-1^PMA^ cells infected or mock infected with RVFV for the indicted times. (F) Immunoblot analysis of DRP1 cleavage in THP-1^PMA^ cells infected or mock infected with RVFV for 2 h, followed by Z-VAD-FMK treatment in the indicted concentration for 22 h. (G) Immunoblot analysis of caspases activation and DRP1 cleavage in THP-1^PMA^ cells infected or mock infected with RVFV for 24 h. Stacked bar graphs show the relative levels of full-length DRP1 and cleaved DRP1 quantified by gray value analysis using ImageJ. Data are representative of three experiments with similar results. For Figure 3: Endogenous DRP1 was detected using an anti-DRP1 monoclonal antibody (CST, #5391).

Furthermore, we found that RVFV infection induced DRP1 cleavage to generate a cleaved DRP1 (∼60 kDa) (Figures 3E). Similar to RVFV infection, apoptotic inducer staurosporine (STS) can also induce DRP1 cleavage (Figures S2B). Cleavage of DRP1 showed a time- and MOI-dependent manner during RVFV infection (Figures 3E and S2C). Exogenously expressed DRP1, either through transient transfection (Figure S2D) or lentiviral transduction (Figure S2E), was also cleaved upon RVFV infection. And Z-VAD-FMK inhibited the formation of cleaved form of DRP1 in a dose-dependent manner (Figure 3F), further supporting that DRP1 might be cleaved by RVFV activated caspases. Indeed, RVFV infection induced the activation of multiple apoptotic caspases, including caspase-3, -6, -7, -8 and -9 (Figure S2F). The results showed that NSs expression alone was sufficient to trigger apoptotic caspases activation and PARP-1 cleavage (Figure S2G). Infection with RVFV^EGFP△NSs^ showed that NSs deletion prevented caspase activation and DRP1 cleavage (Figure 3G, upper panel). Quantification of DRP1 levels confirmed that NSs deletion led to a marked reduction of cleaved DRP1, supporting that NSs is essential for inducing DRP1 cleavage during RVFV infection (Figure 3G, lower panel). These results together suggested that RVFV infection triggered NSs-dependent caspases activation that cleaved DRP1 in infected cells.

### Apoptotic caspase-3, -6, -7 and -8 cleave DRP1 at the AEAD^556^ and D^500^FAD^503^ motifs

To analyze whether activated caspases cleave DRP1, we performed an *in vitro* caspase cleavage assay with purified caspases. Purified DRP1 was incubated with active human recombinant caspase-1, -4, -3, -6, -7, -8 or -9 protein (C1, C4, C3, C6, C7, C8 or C9). Immunoblotting analysis showed that among these caspases, apoptotic caspase-3, -6, -7 and -8, but not caspase-1, -4 or -9, cleaved DRP1 into a C terminal 20 kDa fragment recognized by the anti-strep antibody (Figure 4A). To confirm that caspase-1, -4, -3 and -9 were active under our assay conditions, we performed parallel *in vitro* cleavage assay using the respective substrates. For caspase-1 and -4, incubation with cGAS resulted in a clear reduction in full-length cGAS level, indicating substrate degradation, although the cleavage fragments were not detectable potentially due to antibody epitope affinity (Figure S3A). The caspase-3 cleaved PARP1 (Figure S3B), and caspase-9 cleaved the catalytically inactive mutant procaspase-3 (C3-C163S) (Figure S3C), confirming the activity of these caspases. Pulldown assay showed that DRP1 interacted with caspase-3, -6, -7 and -8 protein, but not with EGFP (Figure 4B). The potential caspase cleavage sites on DRP1 were predicted with the CaspDB database (Kumar et al., 2014). The top five predicted sites were D221, D556, D260, D228 and D503 according to the probability score (Prob.score) (Figure 4C).

**Figure 4.**
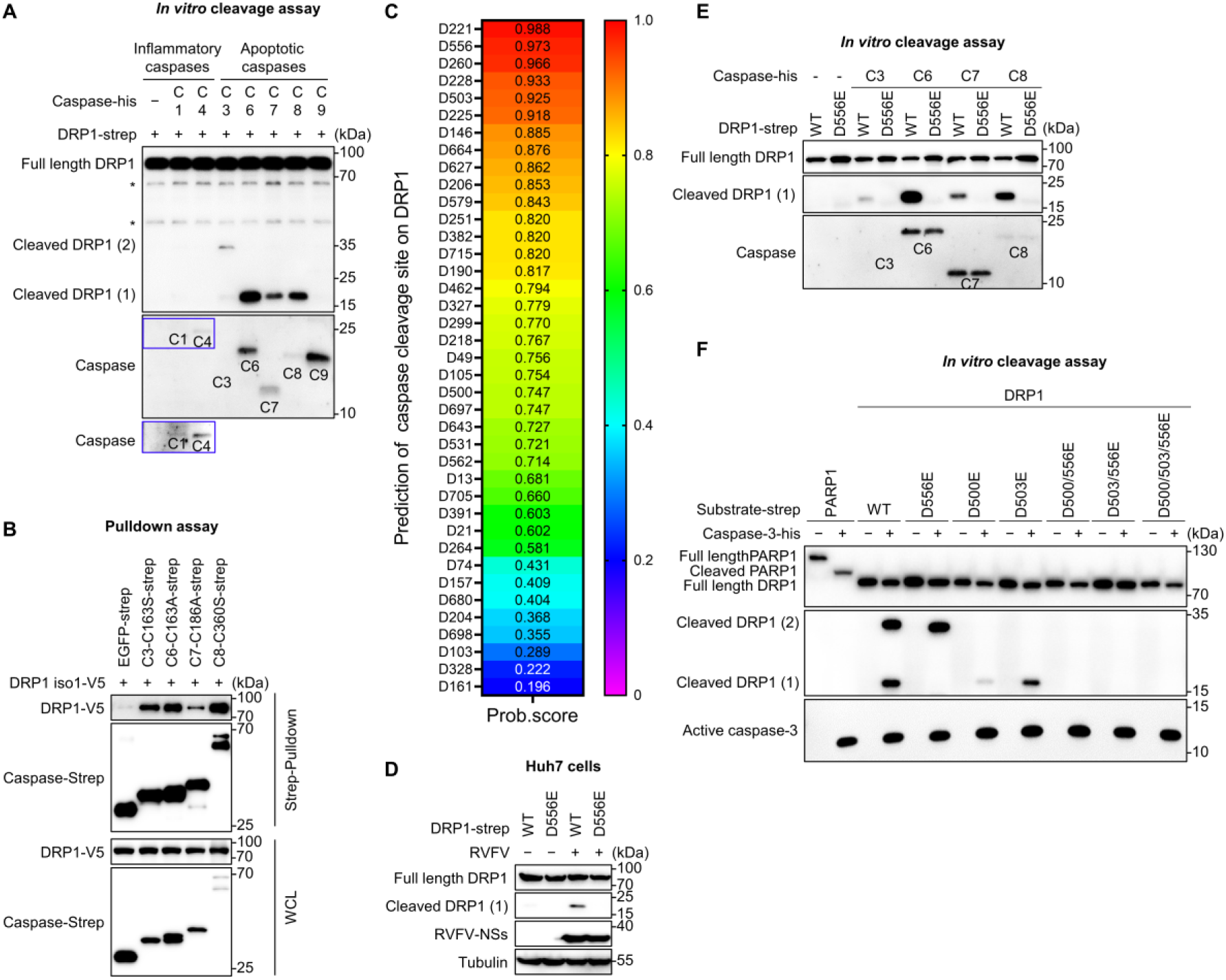
Apoptotic caspase-3, -6, -7 and -8 cleave DRP1 at the AEAD^556^ and/or D^500^FAD^503^ motifs. (A) Immunoblot analysis of DRP1 cleavage by the *in vitro* cleavage assay. The strep-tagged DRP1 protein was expressed and purified from HEK293T cells, and subjected to the *in vitro* cleavage assay with his-tagged caspase-1, 4, 3, 6, 7, 8 or 9 protein, followed by immunoblot analysis. Anti-strep antibody was used to determine the cleavage of DRP1. Anti-His antibody was used to identify caspases. The asterisks (*) represent non-specific bands. The blue solid rectangle represents long exposure. (B) Immunoblot analysis of whole-cell lysates (WCL, bottom) and pull-down proteins with streptavidin magnetic beads (top) from HEK293T cells transfected with indicated plasmids for 24 h. Anti-V5 antibody was used to determine DRP1. Anti-strep antibody was used to identify EGFP or caspases. (C) Prediction of the caspase cleavage sites for DRP1 using the CaspDB database. (D) Immunoblot analysis of DRP1 cleavage in Huh7 cells. Cells were transfected with the DRP1-WT or DRP1-D556E mutant expressing plasmids, followed by RVFV infection or mock infection for 24 h. (E) Immunoblot analysis of DRP1 cleavage by the *in vitro* cleavage assay. The strep-tagged DRP1-WT or DRP1-D556E proteins was expressed and purified from HEK293T cells, and subjected to the *in vitro* cleavage assay as described in (A). (F) Immunoblot analysis of DRP1 cleavage by the *in vitro* cleavage assay. The strep-tagged DRP1-WT or the indicated DRP1 mutants, or his-tagged caspase-3 were expressed and purified from HEK293T cells, and incubated for the *in vitro* cleavage assay followed by immunoblot analysis. DRP1 fragments were detected with anti-strep antibody. Caspase-3 was detected by the anti-His antibody. Data are representative of three experiments with similar results. In Figure 4, exogenous DRP1 was detected with anti-strep antibody (panels A, D, E and F) or anti-V5 antibody (panel B).

In the *in vitro* cleavage assay, caspases cleaved the C-terminal strep tagged DRP1 into two bands that can be recognized by the anti-strep antibody (Figure 4A). One of the cleaved bands was approximately 20 kDa, which corresponds to the potential cleavage site of D556. We therefore substituted the D556 of DRP1 with glutamate (E) to construct the mutant, DRP1-D556E, and transiently expressed the C-terminal strep-tagged DRP1-D556E and DRP1-WT in Huh7 cells. Upon RVFV infection, DRP1-WT was cleaved as detected by the anti-strep antibody, but no corresponding cleaved band was observed in DRP1-D556E expressing cells (Figure 4D), suggesting that D556 may be a key caspase cleavage site of DRP1. The *in vitro* caspase cleavage assay further showed that the DRP1-D556E mutant is resistant to caspase cleavage (Figure 4E), suggesting that caspase-3, -6, -7 and -8 can recognize and cleave DRP1 at the AEAD^556^ motif. Another cleavage band in the caspase-3 treated group was approximately 27 kDa, and there was a canonical caspase recognition motif (D^500^FAD^503^) in this region. We mutated D500 and D503 to glutamate (E) respectively or in combination on the basis of the D556E mutation. The *in vitro* caspase cleavage assay showed that the mutation of either D500 or D503 prevented caspase cleavage and the formation of the 27 kDa fragment (Figure 4F), suggesting that caspase-3 can also recognize and cleave DRP1 at the D^500^FAD^503^ motif and both D residues are important for this cleavage. Thus, apoptotic caspase-3, -6, -7 and -8 cleave DRP1 at the AEAD^556^ motif and caspase-3 has an additional cleavage site at the D^500^FAD^503^ motif (Figure S3D).

### Caspases-mediated DRP1 cleavage promotes anti-viral innate immunity

Mitochondria is a critical platform for regulating innate immune responses (Arnoult et al., 2011), so we reasoned that RVFV infection triggered DRP1 down-regulation and mitochondrial elongation may regulate anti-viral immune responses. To analyze this possibility, we first depleted DRP1 in THP-1 cells through knockdown (KD) by shRNA or knockout (KO) by sgRNA with CRISPR-Cas9. The RVFV-induced transcription of IFNβ and its downstream interferon-stimulated genes (ISGs), including RSAD2, CXCL10, ISG15, ISG54 and ISG56, were significantly enhanced in DRP1 knockout macrophages compared with the control cells (Figure 5A). Immunoblot analysis showed that DRP1 depletion through knockdown (Figure S4A) or knockout (Figure 5B) enhanced the phosphorylation of TBK1 and up-regulated the expression of ISGs after RVFV infection indicating elevated innate immune responses. In line with this, intracellular RVFV-NP protein expression (Figures S4A and 5B), or RVFV-NSs protein expression (Figure S4C) and infectious RVFV production (Figures 5C and S4B) were all significantly reduced in DRP1-depleted macrophages relative to the control cells. Treatment with DRP1 inhibitor mdivi-1, which inhibits DRP1 fission functionality, also inhibited RVFV replication in a dose-dependent manner (Figure S4D). Restriction of RVFV replication was also observed in mouse embryonic fibroblast (MEF) cells with DRP1 depletion by CRISPR-Cas9 mediated knockout (Figure S4E). Additionally, we observed that DRP1 depletion induced mitochondrial elongation in cells (Figure 5D). These results suggest that the altered mitochondrial morphology may underlie the potentiated innate antiviral responses resulting from DRP1 depletion.

**Figure 5.**
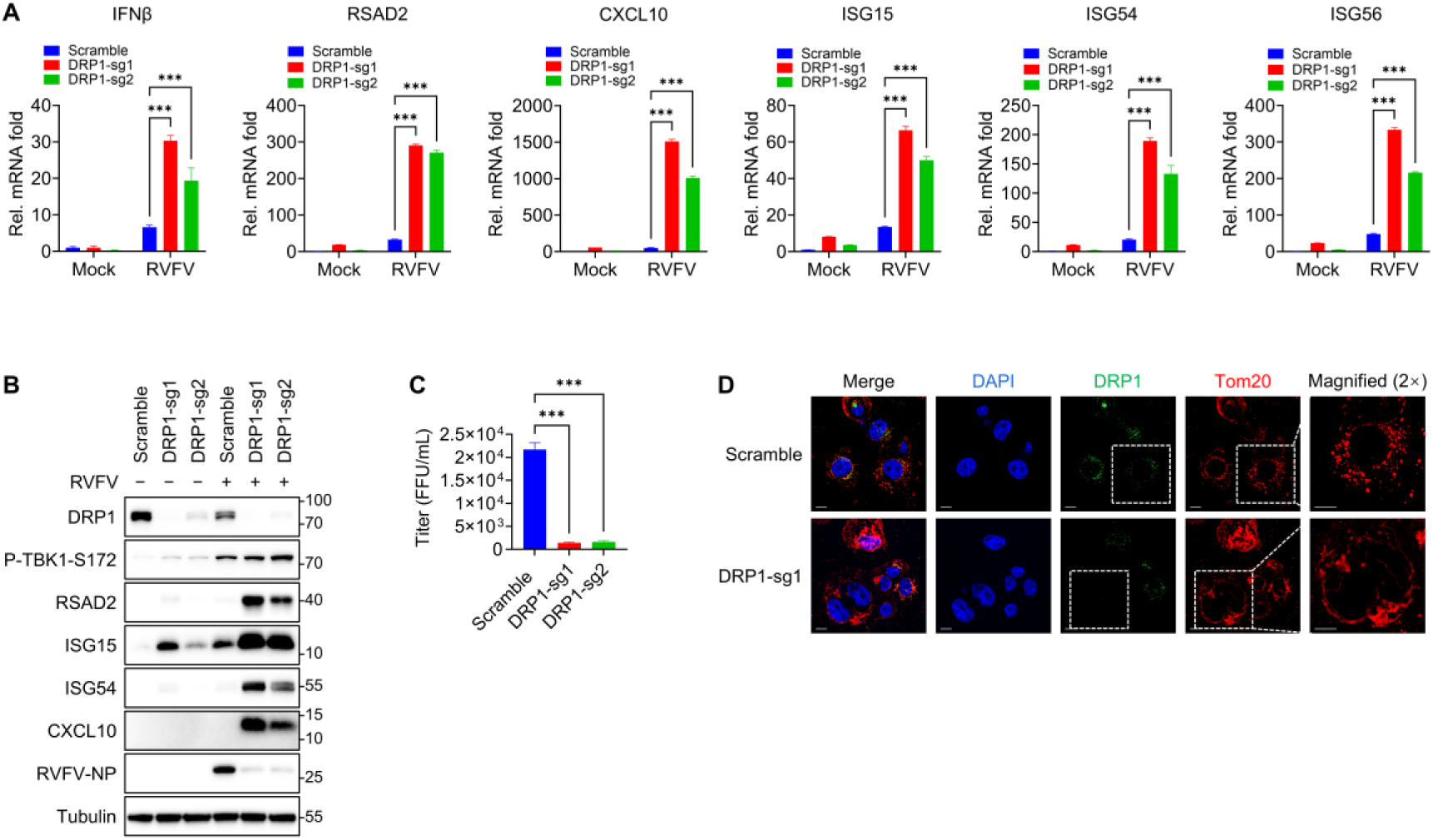
DRP1 deficiency promotes antiviral innate immunity. (A) qPCR analysis of the indicated genes in scramble or DRP1 knockout THP-1^PMA^ cells infected or mock infected with RVFV for 12 h. \*\*\**P* < 0.001 (Two-way ANOVA test). (B) Immunoblot analysis of the indicated proteins in scramble or DRP1 knockout THP-1^PMA^ cells infected or mock infected with RVFV for 24h. (C) Focus forming assay to determine the titer from RVFV infected scramble or DRP1 knockout THP-1^PMA^ cells infected for 24h. \*\*\**P* < 0.001 (One-way ANOVA test). (D) Immunofluorescence analysis of mitochondrial morphology in Scramble and DRP1 knockout THP-1^PMA^ cells. Mitochondria were labeled with anti-Tom20 antibody (red), DRP1 was labeled with anti-DRP1 antibody (green, # 8570, CST) and Nuclei were stained with DAPI (blue). Scale bars, 10 μm.Data are representative of three experiments with similar results.

In contrast, overexpression of DRP1 inhibited the up-regulation of IFNβ and its downstream representative ISGs (RSAD2, CXCL10, ISG15, ISG54 and ISG56) mRNA triggered by RVFV infection compared with the control cells (Figures 6A). DRP1 overexpression also reduced the phosphorylation of TBK1 and inhibited the up-regulation of ISGs (RSAD2, ISG15 and ISG54) after RVFV infection (Figure 6B). Consistently, intracellular RVFV-NP protein level (Figure 6B) and infectious RVFV production (Figure 6C) were significantly higher in DRP1-overexpressed macrophages relative to the control cells. These results suggested that over-expression of DRP1 dampened innate antiviral responses.

**Figure 6.**
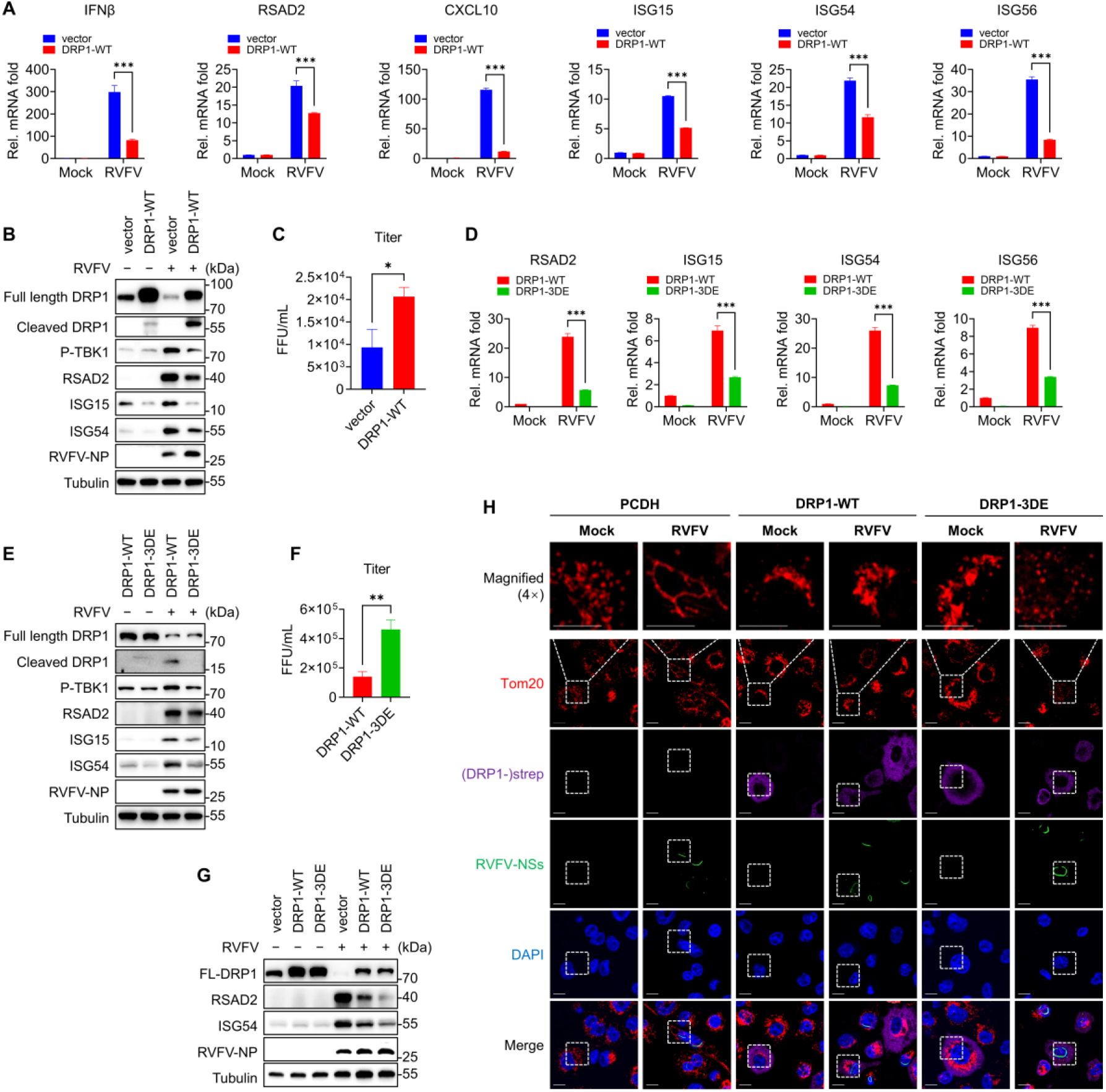
Caspases-mediated DRP1 cleavage promotes antiviral innate immunity. (A) qPCR analysis of the indicated genes in vector or DRP1-WT transduced THP-1^PMA^ cells infected or mock infected with RVFV for 12 h. \*\*\**P* < 0.001 (Two-way ANOVA test). (B) Immunoblot analysis of the indicated proteins in vector or DRP1-WT transduced THP-1^PMA^ cells infected or mock infected with RVFV for 24h. (C) Focus forming assay to determine the titer from RVFV infected vector or DRP1-WT transduced THP-1^PMA^ cells infected with RVFV for 24h. \**P* < 0.05, \*\**P* < 0.01, \*\*\**P* < 0.001 (Student’s t test). (D) qPCR analysis of the indicated genes in DRP1-WT or DRP1-3DE transduced THP-1^PMA^ cells infected or mock infected with RVFV for 12 h. \*\*\**P* < 0.001 (Two-way ANOVA test). (E) Immunoblot analysis of the indicated proteins in DRP1-WT or DRP1-3DE transduced THP-1^PMA^ cells infected or mock infected with RVFV for 24 h. (F) Focus forming assay to determine the titer from RVFV infected DRP1-WT or DRP1-3DE transduced THP-1^PMA^ cells infected with RVFV for 24 h. \*\**P* < 0.01 (Student’s t test). (G) Immunoblot analysis of the indicated proteins in vector, DRP1-WT or DRP1-3DE transduced THP-1^PMA^ cells infected or mock infected with RVFV for 24 h. (H) Immunofluorescence analysis of mitochondrial morphology in PCDH (control), DRP1-WT, or DRP1-3DE transduced THP-1^PMA^ cells, infected or mock infected with RVFV for 24 h. Mitochondria were labeled with anti-Tom20 antibody (red), DRP1-WT and DRP1-3DE were labeled with anti-strep antibody (purple), RVFV were labeled with anti-RVFV-NSs antibody (green) and Nuclei were stained with DAPI (blue). Scale bars, 10 μm. Data are representative of three experiments with similar results.

To verify whether DRP1 cleavage by apoptotic caspases play a role in regulating immune responses, human DRP1-WT or caspase-resistant DRP1-3DE (a mutant of which the three D residues in the caspase cleavage motifs D^500^FAD^503^ and AEAD^556^ were substituted into E) was over-expressed in THP-1 cells. We found that cells expressing DRP1-3DE, which is resistant to caspase cleavage, showed further attenuated transcription of ISGs RSAD2, ISG15, ISG54 and ISG56 mRNA compared with the DRP1-WT over-expressing cells, in response to RVFV infection (Figure 6D). Compared with DRP1-WT, DRP1-3DE also more strongly inhibited the phosphorylation of TBK1 and the up-regulation of ISGs after RVFV infection (Figures 6E and 6G). Additionally, intracellular RVFV-NP protein expression (Figures 6E and 6G) and infectious RVFV production (Figure 6F) were significantly higher in DRP1-3DE expressing macrophages relative to the DRP1-WT expressing cells. To examine the impact of these DRP1 variants on mitochondrial morphology during RVFV infection, we analyzed these cells by immunofluorescence. Control (empty vector-transduced) THP-1^PMA^ cells exhibited mitochondrial elongation upon RVFV infection, whereas overexpression of either DRP1-WT or DRP1-3DE prevented RVFV-induced mitochondrial elongation (Figure 6H). These results underscore the role of DRP1 in maintaining mitochondrial morphology.

Together, these data revealed that while both DRP1-WT and the cleavage-resistant DRP1-3DE maintain mitochondrial fragmentation, the DRP1-3DE mutant showed a more potent inhibition of antiviral immune responses. This indicates that caspases-mediated DRP1 cleavage plays a role in up-regulating anti-viral innate immune responses.

### Caspases-mediated DRP1 cleavage leads to mitochondrial elongation and promotes MAVS aggregation

The above evidences suggest that the DRP1-depletion altered mitochondrial morphodynamics is linked with regulation of antiviral innate immunity. Activation and aggregation of the mitochondria-associated MAVS is a key event to mediate the activation of TBK1/IKKε and the down-stream immune responses (Chen et al., 2020; Zhang et al., 2020). We therefore analyzed whether mitochondrial elongation caused by caspases-mediated DRP1 cleavage can facilitate MAVS aggregation to promote the antiviral innate immunity. Confocal fluorescence microscopy revealed that DRP1 depletion in macrophages caused significant MAVS aggregation on fused mitochondria (Figure 7A). Concomitantly, DRP1 deficiency in THP-1^PMA^ cells either through CRISPR-Cas9 knockout showed significantly increased transcription of representative ISGs including RSAD2, ISG56 and CXCL10 mRNA compared with the scramble cells (Figure S5A). Similarly, DRP1 deficiency in MEF cells through knockout also elevated transcription of representative ISGs RSAD2, ISG56 and CXCL10 mRNA compared with the scramble cells (Figure S5B). RVFV infection of THP-1^PMA^ led to caspase activation, DRP1 depletion and aggregation of MAVS (Figure 7B). DRP1 depletion further increased aggregation of MAVS upon SeV (Figure 7C) or RVFV infection (Figure 7D). Simultaneous knock-out (dKO) of DRP1 and MAVS strongly attenuated the elevated antiviral immune responses observed in RVFV infected DRP1 KO cells and promoted viral replication (Figures 7E and 7F). These data suggest that DRP1 depletion promotes antiviral innate immunity, at least in part, through inducing MAVS aggregation and activation. Altogether, these data indicated that virus infection can trigger caspases-mediated DRP1 cleavage leading to mitochondrial elongation, which enhanced MAVS aggregation to elevate the antiviral immune responses.

**Figure 7.**
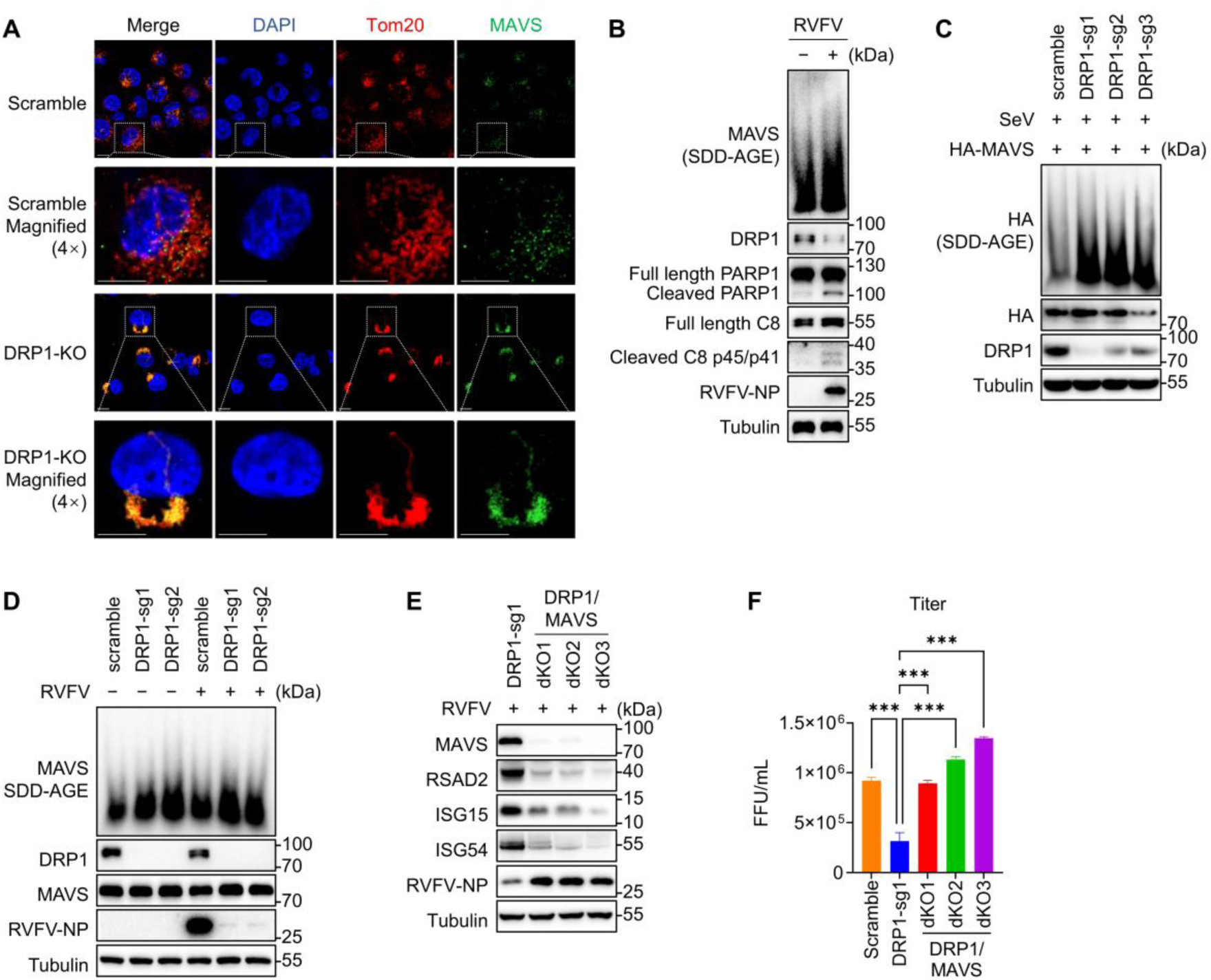
Caspases-mediated DRP1 cleavage induces mitochondrial elongation to promote MAVS aggregation and immune responses. (A) Immunofluorescence analysis of MAVS aggregation in scramble or DRP1 knockout THP-1^PMA^ cells. Mitochondria were labeled with anti-Tom20 antibody (red), MAVS was labeled with anti-MAVS antibody (green) and Nuclei were stained with DAPI (blue). Scale bars, 10 μm. (B) Immunoblot analysis of caspase-8 activation, DRP1 protein level and MAVS aggregation in THP-1^PMA^ cells infected or mock infected with RVFV or 24 h. (C) Immunoblot analysis of MAVS aggregation in scramble or DRP1 knockout HEK293T cells. HEK293T cells were transfected with HA-tagged MAVS (HA-MAVS) plasmid for 24 h, followed by SeV infection for 12 h. (D) Immunoblot analysis of MAVS aggregation in scramble or DRP1 knockout THP-1^PMA^ cells infected or mock infected with RVFV for 24 h. (E) Immunoblot analysis of the indicated proteins in DRP1 knockout or DRP1 and MAVS double knockout THP-1^PMA^ cells infected with RVFV for 24 h. (F) Focus forming assay to determine the titer from RVFV infected DRP1 knockout or DRP1 and MAVS double knockout THP-1^PMA^ cells infected for 24 h. \*\*\**P* < 0.001 (One-way ANOVA test). Data are representative of three experiments with similar results.

### Caspases-mediated DRP1 cleavage and elevation of immune responses is conserved among different viruses infection

Next, we analyzed whether caspase-dependent DRP1 cleavage and promotion of innate immune responses is a general mechanism during virus infection. For this purpose, H1N1 and SeV were selected as additional RNA model viruses, and HSV-1 was selected as a DNA model virus. THP-1^PMA^ cells were infected with H1N1, SeV or HSV-1 and the activation of multiple apoptotic caspases, including caspase-3, -6, -7, -8 and -9 were observed (Figures 8A-8C). Concomitantly with caspases activation, the cleavage of DRP1 was observed in H1N1, SeV or HSV-1 infected cells (Figures 8A-8C). Mitochondrial elongation was also observed in H1N1, SeV or HSV-1 infected cells (Figure 8D). Similar with RVFV, H1N1, SeV or HSV-1 infection induced DRP1 depletion was prevented by Z-VAD-FMK treatment (Figures 8E-8G). Additionally, infectious H1N1 or HSV-1 titers were all significantly reduced in DRP1-depleted macrophages relative to the control cells (Figures 8H and 8I), suggesting that DRP1 depletion may pose restriction effect on H1N1 and HSV-1 replication, similar with the case of RVFV infection. Whether and how this caspase-mediated DRP1 cleavage per se affects progeny virus titers requires further investigation. These results suggested that the caspase-triggered DRP1 cleavage and elevation of immune responses is a general mechanism occurring during virus infection (Figure 9).

**Figure 8.**
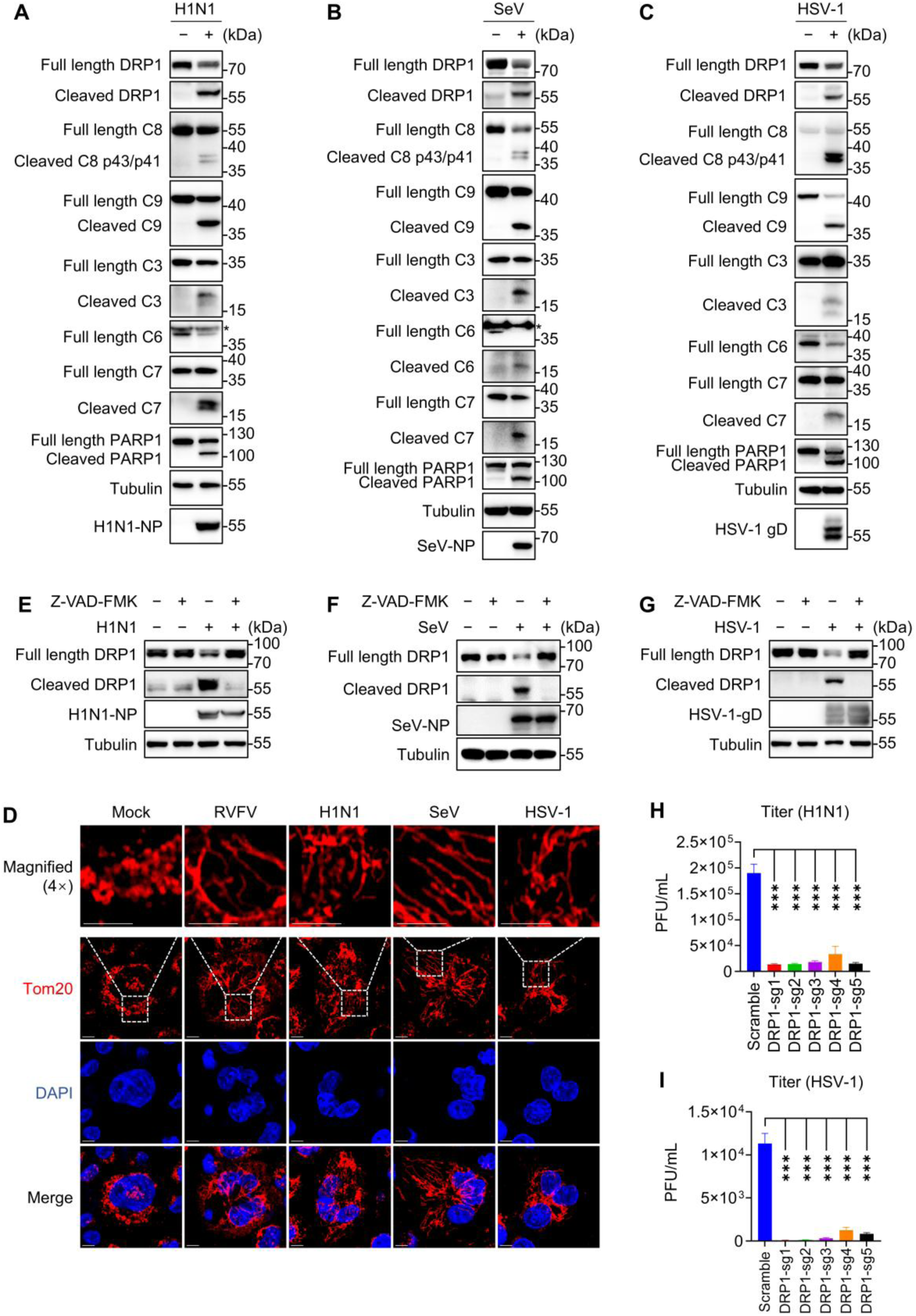
Multiple viruses trigger caspases-mediated DRP1 cleavage. (A-C) Immunoblot analysis of caspases activation and DRP1 cleavage in THP-1^PMA^ cells infected or mock infected with H1N1 (A), SeV (B) or HSV-1 (C) for 24 h. The asterisks (*) represent non-specific bands. (D) Mitochondria were visualized by immunofluorescence using confocal microscopy. THP-1^PMA^ cells were infected or mock infected with RVFV, H1N1, SeV or HSV-1 for 18 h, respectively. Mitochondria were labeled with anti-Tom20 (red) antibody and Nuclei were stained with DAPI (blue). Scale bars, 7.5 μm.(E) Immunoblot analysis of DRP1 cleavage in THP-1^PMA^ cells infected or mock infected with H1N1 for 2 h, followed by Z-VAD-FMK treatment for 22 h. (F) Immunoblot analysis of DRP1 cleavage in THP-1^PMA^ cells infected or mock infected with SeV for 2 h, followed by Z-VAD-FMK treatment for 10 h. (G) Immunoblot analysis of DRP1 cleavage in THP-1^PMA^ cells infected or mock infected with HSV-1 for 2 h, followed by Z-VAD-FMK treatment for 22 h. (H-I) Plaque assay results determining the titers from virus infected scramble or DRP1 knockout THP-1^PMA^ cells infected with H1N1 (H) or HSV-1 (I) for 24 h. \*\*\**P* < 0.001 (One-way ANOVA test). Data are representative of three experiments with similar results.

**Figure 9.**
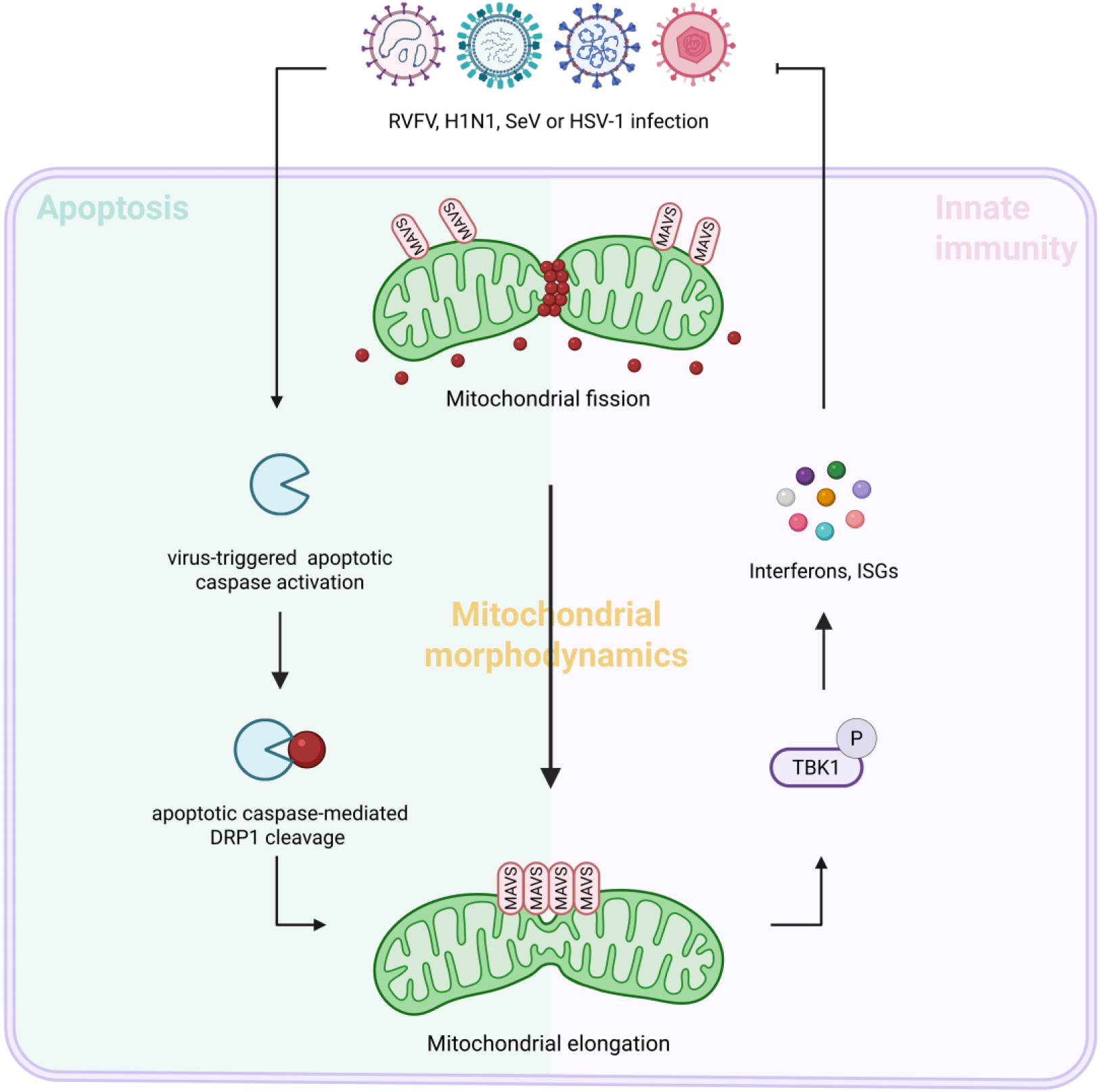
Model of apoptotic caspases-mediated DRP1 cleavage enhances anti-viral innate immunity. **Related to Figure 8.** Viral infection triggers the activation of apoptotic caspases, including caspase-3, -6, -7 and 8 (C3, C6, C7 and C8). Activated apoptotic caspases cleave and inactive DRP1 at the AEAD^556^ and/or D^500^FAD^503^ motifs and induces mitochondrial elongation. Elongated mitochondria promote MAVS aggregation and elevate anti-viral immune responses.

## Discussion

Virus triggered apoptotic caspase activation has been shown to dampen antiviral innate immunity via the cleavage of multiple key immune molecules, such as cGAS, MAVS, IRF3, and NF-κB members (Ning et al., 2019; Wu et al., 2022). Whether apoptotic caspases can promote innate immune responses to inhibit viral infection is currently unknown. Here, we demonstrated that virus-activated apoptotic caspases, including caspase-3, -6, -7 and -8, cleave and deplete the mitochondrial fission factor DRP1, leading to mitochondrial elongation, which in turn results in increased MAVS aggregation and enhanced antiviral innate immune signaling. This mechanism, observed during infection with multiple RNA and DNA viruses, reveals a previously unrecognized role for apoptotic caspases in promoting innate immunity. Consequently, the net immunological effect of caspase activation is not universally inhibitory but is pathway-and context-dependent. Together with the established role of caspase-mediated cell death in eliminating viral replication niches, these findings indicate that apoptotic caspase is co-opted by the host to exert a dual antiviral action: by cleaving DRP1 to enhance mitochondrial signaling, and by eliminating infected cells through programmed death.

The dynamic balance between fission and fusion plays a critical role in maintaining the functionality of mitochondria. DRP1 is the major regulator of mitochondrial fission, and its activity is regulated by multiple post-translational modifications, including phosphorylation, ubiquitination, sumoylation, and S-nitrosylation (Khan et al., 2015). Previous reports showed that viruses can dampen the activity of DRP1 by modulating its phosphorylation state. For example, SeV infection-induced TBK1-mediated DRP1 phosphorylation and abrogates its mitochondrial fission function, leading to mitochondrial elongation (Chen et al., 2020). Moreover, DENV NS4B inhibits phospho-S616-dependent activation of DRP1 and its subsequent translocation to mitochondria, thus inducing mitochondrial elongation in virus-infected cells (Chatel-Chaix et al., 2016). Here, we discovered that cleavage by multiple caspases is a novel mechanism to dampen DRP1 functionality and enhance innate immune responses. This new layer of regulation adds to the complex regulation mechanisms of DRP1 and further indicates its critical role in maintaining mitochondrial morphodynamics and immune responses.

It is found that multiple caspases, including caspase-3, -6, -7 and -8, are capable of cleaving DRP1. Among them, caspase-8 is the initiator apoptotic caspase, while caspase-3, -6 and -7 belong to the executive apoptotic caspases. In addition, multiple sites in DRP1 are sensitive to caspase cleavage, including the D^500^FAD^503^ and AEAD^556^ motifs. The susceptibility of DRP1 to cleavage by different types of caspases and the presence of multiple caspase-sensitive sites in DRP1 ensure the efficient cleavage of DRP1 under different contexts. Indeed, multiple viruses and other pathogens trigger apoptotic caspases activation during infection through diverse mechanisms (Fang and Peng, 2021; Li et al., 2020b). The results that, besides RVFV, H1N1, SeV and HSV-1 infection triggered caspase-dependent cleavage of DRP1 further support the notion. It can be speculated that DRP1 potentially serves as a common sensor of pathogen invasion through sensing pathogen activated caspases and promotes innate immunity. This might be a general host defense mechanism occurring under different pathogen invasion contexts.

Our *in vitro* findings indicates the caspase–DRP1 axis as an antiviral mechanism, yet its *in vivo* validation needs to consider the complex biology of DRP1. This key fission factor is regulated by several post-translational modifications. Given the embryonic lethality of constitutive Drp1 knockout, tissue-specific conditional knockout models may be used to delineate its role in the immune system *in vivo*. We anticipate that such models would provide evidences regarding the physiological relevance of mitochondrial dynamics in antiviral defense.

In summary, these results revealed an apoptotic caspase-DRP1 axis in promoting anti-viral immune responses through regulating mitochondrial morphodynamics. This indicates that apoptotic caspases activation can have both pro-or anti-viral effects depending on the cleaved substrates. During the course of virus infection, it is likely that the balance between pro-or anti-viral effects is controlled by multiple viral and host factors. The overall effect of how caspase activation regulates viral infection and pathogenesis may be context-dependent and awaits further characterization for different viruses.

### Limitation

A limitation of this study is that the overexpression system used to compare DRP1-WT and DRP1-3DE abundance may not fully recapitulate endogenous DRP1 regulation. In this system, RVFV infection reduced the full-length DRP1 level of both DRP1-WT and DRP1-3DE to a similar extent (Figure 6E), suggesting that additional caspase-independent mechanisms may contribute to regulation of total DRP1 abundance in this overexpression system. This may be associated with the strong transcription inhibition effect of RVFV NSs protein. Future studies using endogenous DRP1-3DE knock-in models and *in vivo* infection models will help clarify this issue and strengthen the biological significance.

## Materials and Methods

### Plasmids

cDNAs encoding human caspase-3 (CCDS3836.1) was synthesized by Tsingke (Wuhan, China) and cloned into pRK plasmid with a C-terminal His tag. Plasmids, including C-terminal strep-tagged human DRP1 (hDRP1) and viral proteins of RVFV (NSs, NP, NSm, Gn, Gc and Pol), were constructed by standard molecular biology techniques. Point mutants were generated by using the ClonExpress II One Step Cloning Kit (#C112-01, Vazyme, China) and the construct coding the wild-type DRP1 as template. Each construct was confirmed by sequencing.

### Antibodies, recombinant proteins and reagents

The anti-V5 (#V8012) antibody was purchased from Sigma (St. Louis, MO, USA). The anti-strep (#A00626, pAb for Western blot; #A01732, mAb for immunofluorescence analysis) antibody was purchased from GenScript (Nanjing, China). The anti-MAVS (#sc-166583) antibody used in immunofluorescence analysis was purchased from Santa Cruz Biotechnology (Delaware, USA). The anti-Tom20 (#ab209606) was purchased from Abcam (Cambridge, UK). The anti-DRP1 (#5391 for Western blot; # 8570 for immunofluorescence analysis), anti-caspase-3 (#9662), anti-cleaved caspase-3 (#9661), anti-caspase-6 (#9762), anti-cleaved caspase-6 (#9761), anti-caspase-7 (#9492), anti-caspase-8 (#9746), anti-cleaved caspase-8 (#9496), anti-caspase-9 (#9502), anti-PARP1 (#9542), anti-MAVS (#24930), anti-TBK1 (#38066) and anti-Phospho-TBK1/NAK (Ser172) (#5483) antibodies were obtained from Cell Signaling Technology (Beverly, MA, USA). The anti-MFN2 (#A10175) and anti-OPA1 (#A9833) antibodies were obtained from ABclonal. The anti-His tag (#66005-1-Ig), anti-MFN1 (#13798-1-AP), anti-RSAD2 (#28089-1-AP), anti-MAVS (#14341-1-AP) used in immunoblot analysis, anti-alpha Tubulin (#14555-1-AP), anti-GAPDH (#60004-1-Ig), goat anti-mouse (#SA00001-1) and goat anti-rabbit (#SA00001-2) IgG-horseradish peroxidase (HRP) secondary antibodies were purchased from Proteintech (Wuhan, China). Anti-Sendai Virus pAb (#PD029) was purchased from Medical & Biological Laboratories (Japan). The rabbit anti-RVFV-NP and rabbit anti-RVFV-NSs antibodies were made in-house. Alexa Fluor 488 goat anti-mouse IgG (H + L) (#A11029) and Alexa Fluor 568 goat anti-rabbit IgG (H + L) (#A11036) were purchased from Thermo Fisher.

Active human recombinant caspase-4 (#1084) and caspase-7 (#1087-25) proteins were purchased from Biovision. Active human recombinant caspase-1 (#268-10011-1), caspase-6 (#268-10016-1) and caspase-8 (#268-10018-1) proteins were obtained from Raybiotech. Active human recombinant caspase-3 (#CC119) and caspase-9 (#CC120) proteins were purchased from Merck.

PMA (#P1585) was purchased from Sigma-Aldrich (St. Louis, MO, USA). Recombinant murine M-CSF (#315-02) was obtained from PeproTech. MG132 (#S2619), Chloroquine diphosphate (#S4157), Z-VAD-FMK (#S7023) and Mdivi-1 (#S7162) were obtained from Selleck (Houston, TX, USA). Staurosporine (#HY-15141) was purchased from MedChemExpress (New Jersey, USA). DAPI (#C1002) was purchased from Beyotime (Shanghai, China).

### Cell lines

The HEK293T, BHK-21, Vero, A549, and THP-1 cell lines were purchased from American Type Culture Collection (ATCC). Huh7 cell was obtained from China Center for Type Culture Collection and the MDCK cell line was provided by Dr Jianjun Chen (Wuhan institute of virology, China). Adherent cells were cultured in Dulbecco’s modified Eagle’s medium (DMEM; Gibco) supplied with 10% fetal bovine serum (FBS; Gibco) and 1% Pen Strep antibiotics (Gibco), THP-1cells were cultured in RPMI 1640 (Gibco) containing 10% FBS (Gibco) and 1% Pen Strep (Gibco). All cells were cultured at 37°C in a humidified 5% CO_2_ incubator.

### Viruses and titer determination

Recombinant wild-type RVFV (RVFV) and the NSs deleted RVFV (RVFV^EGFP△NSs^) were rescued by T7 RNA polymerase-driven reverse genetics system as previously described (Li et al., 2019). And RVFV strains were propagated in BHK-21 cells under Biosafety Level-3 (BSL-3) conditions. All RVFV-related experiments were performed at the BSL-3 laboratory of the Wuhan Institute of Virology, Chinese Academy of Sciences (Wuhan, China) according to the institutional guidelines.

Titer of RVFV was determined by focus forming assay on Vero cells as previously described (Li et al., 2020a). Briefly, confluent Vero monolayers were infected with 10-fold dilutions of virus and incubated at 37°C for 1 h, then culture medium was replaced by DMEM containing 2% FBS and 1.1% (wt/vol) carboxymethyl-cellulose/ methylcellulose. After 3 days of incubation, cells were fixed with 3.6%-4% formaldehyde. And then, Foci were visualized by DAB staining after two-step immunostaining with an antibody against viral protein NP and an anti-rabbit horseradish peroxidase-conjugated secondary antibody.

Plaque assays were performed on Vero cells or MDCK cells to determine the titer of HSV-1 or H1N1, respectively. For HSV-1, confluent Vero monolayers were infected with 10-fold dilutions of virus and incubated at 37°C for 1 h, then culture medium was replaced by DMEM containing 2% FBS. After 48 h of incubation, cells were stained with 0.5% crystal violet after fixing with 3.6%-4% formaldehyde and plaques were counted. For H1N1, we referred to the protocol of Galani et al (Galani et al., 2022).

### Liver samples of mice

BALB/c mice were purchased from Charles River Laboratories (Beijing, China). 6-week-old BALB/c mice were intraperitoneally injected or mock injected with 1000 plaque forming unit (PFU) of RVFV in 100 µL of PBS. Mice were monitored for 3 days and Liver samples were harvested for Immunoblot analysis. Animal experiments were performed in agreement with Regulations for the Administration of Affairs Concerning Experimental Animals in China. The procedures were approved by the Laboratory Animal Care and Use Committee of the Wuhan Institute of Virology, Chinese Academy of Sciences (Wuhan, China). The approval code was WIVA38202002.

### Cell differentiation

THP-1 cells were differentiated into macrophages by treatment with 40 ng/mL of PMA in RPMI 1640 (Gibco) containing 10% FBS (Gibco) and 1% Pen Strep (Gibco) at 37°C for 24 h, and then cells were cultured in RPMI 1640 (Gibco) containing 10% FBS (Gibco) and 1% Pen Strep (Gibco) without PMA at 37°C for 24 h.

### Generation of BMDMs

Bone marrow cells were isolated from tibiae and femur of mice. The bone marrow cells were cultured in RPMI 1640 medium containing 10% FBS, 1% Pen Strep and 20ng/ml M-CSF at 37°C in a humidified 5% CO_2_ incubator for 3-5 days to generation of BMDMs.

### Knockdown, knockout, double-knockout and overexpression

Knockdown of DRP1 with shRNA was done by lentiviral transduction of THP-1 cells. Knockout of DRP1 or double Knockout of DRP1 and MAVS with sgRNA and Cas9 was accomplished by transduction of THP-1 cells. Overexpression of DRP1-WT or DRP1 mutant with coding sequence was done by lentiviral transduction of THP-1 cells. Sequences of targeting shRNAs and sgRNAs used in this study were listed in Table S1.

### Fluorescence and Immunofluorescence Microscopy

THP-1^PMA^ cells, Huh7 cells or A549 cells were infected or mock infected with RVFV, and then fixed with 4% paraformaldehyde (PFA) at room temperature for 30 min, permeabilized with 0.3% (vol/vol) Triton X-100 for 20 min, blocked in 3% bovine serum albumin in PBS for 30 min, and incubated sequentially with the indicted primary antibodies at room temperature for 1h, followed by incubation with the correspondingly fluorescent-dye conjugated secondary antibodies. Cells were stained with DAPI, mounted with ProLongTM Gold antifade reagent (Invitrogen, San Diego, CA, USA), and then analyzed using confocal microscope (Andor Dragonfly 202). Images were processed by Imaris.

### Transmission Electron Microscope (TEM)

THP-1^PMA^ cells were infected or mock infected with RVFV at MOI = 1 for 30 h. Cells were fixed with 2.5% (w/v) glutaraldehyde in 0.1 M sodium chloride and and processed for Electron Microscopy (EM). The mitochondria were observed under a transmission electron microscope (FEI Tecnai G2 microscope at 200 kV).

### Pulldown assay

HEK293T cells were transfected with the indicated plasmids for 24 h. The cells were washed twice with PBS, lysed with RIPA (weak) buffer (#P0013D, Beyotime, Shanghai, China) and rotated for 30 min at 4°C. After centrifugation at 16000× *g* for 20 min, the whole cell lysates were incubated with streptavidin magnetic beads (#2-4090-010, Iba-Lifesciences, Germany) and rotated overnight at 4°C on a rotator. After washing three times with PBST, the pull-down complexes were eluted with 2× SDS sample loading buffer for 10 min at 95°C and applied to immunoblot analysis.

### Immunoblot Analysis

Cells were lysed with buffer (#P0013, Beyotime, Shanghai, China). Cell lysates were subjected to SDS-polyacrylamide gel electrophoresis (PAGE) then transferred to polyvinylidene difluoride (PVDF) membranes (Millipore, Billerica, MA, USA). Proteins were incubated with primary antibodies, then secondary horseradish peroxidase-conjugated goat anti-rabbit/mouse IgG. Protein bands were detected by an enhanced chemiluminescence (ECL) kit (Millipore, Billerica, MA, USA) using a Chemiluminescence Analyzer (Chemiscope600pro, Shanghai, China). The gray values of the Western blot bands were analyzed using ImageJ software (National Institutes of Health, USA).

### *In vitro* cleavage assay

*In vitro* protease cleavage experiments were performed as previously described (Wang et al., 2017). Briefly, recombinant or purified caspase was incubated with DRP1-WT or DRP1 mutant in cleavage buffer (100 mM HEPES, 10% (w/v) sucrose, 0.1% (w/v) CHAPS, PH 7.0, 10mM DTT) at 37°C for 2 h and then samples were applied to immunoblot analysis.

### Semi-denaturing detergent agarose gel electrophoresis (SDD-AGE)

Cells were lysed with NP-40 buffer (#P0013F, Beyotime, Shanghai, China) and mixed with 5× sample buffer (2.5× TBE, 50% glycerol, 10% SDS and 0.0125% bromophenol blue). The samples were loaded onto a 1.5% agarose gel in 1× TBE containing a final concentration of 0.1% SDS, electrophoresed at a constant voltage of 100 V at 4°C until the bromophenol blue dye reached the bottom of the agarose gel, and then transferred to PVDF membranes for immunoblot analysis.

### RNA isolation and Quantitative RT-PCR analysis

Total RNAs were extracted with RNAiso Plus (Takara, Japan) following the manufacturer’s instructions. qRT-PCR was performed with a two-step procedure using the HiScript II 1st Strand cDNA Synthesis Kit (+gDNA wiper) (#R212-01, Vazyme, China) and ChamQ Universal SYBR qPCR Master Mix (#Q711-02/03, Vazyme, China). Data shown are the relative abundance of the indicated mRNA normalized to that of GAPDH. Primer sequences are available upon request. Gene-specific primer sequences were listed in Table S2.

### Statistical analysis

All data are presented as mean values ± SD of three or more experiments. Statistical significance was determined with Student’s t test, One-way ANOVA or Two-way ANOVA as showed in figure legends with *P* < 0.05 considered statistically significant. ns, not significant; \**P* < 0.05, \*\**P* < 0.01, \*\*\**P* < 0.001.

### Software

Data were analyzed with GraghPad prism 9. Images were processed by Imaris or ImageJ.

## Acknowledgments

This work was supported by the Prevention and Control of Emerging and Major Infectious Diseases-National Science and Technology Major Project (Grant 2026ZD01910600), Major Project of Guangzhou National Laboratory, Grant No.SRPG22-001/Prevention and Control of Emerging and Major Infectious Diseases-National Science and Technology Major Project, Grant No. 2026ZD01999700, National Natural Science Foundation of China (numbers 82341082, 32070179), Fundamental Research Program of Shanxi Province (No.202403021222310), Scientific and Technologial Innovation Programs of Higher Education Institutions in Shanxi (No.2024L282), and Changzhi Medical College Doctoral Research Startup Fund Project (No. 2024BS21). We thank Dr. Qinxue Hu (Wuhan Institute of Virology), Dr. Jianjun Chen (Wuhan Institute of Virology) and Dr. Yuchen Xia (Wuhan University) for help with materials, and Dr. Leike Zhang (Wuhan Institute of Virology) and Dr. Peng Zhou (Wuhan Institute of Virology) for their advice and guidance. We thank Hao Tang, Jun Liu and Jia Wu from BSL-3 Laboratory of Wuhan Institute of Virology for their critical support. We thank Ding Gao and Pei Zhang from Center for Instrumental Analysis and Metrology at Wuhan Institute of Virology for their help with confocal microscopy and transmission electron microscope experiments.

## Supplemental Information

**Figure S1.**
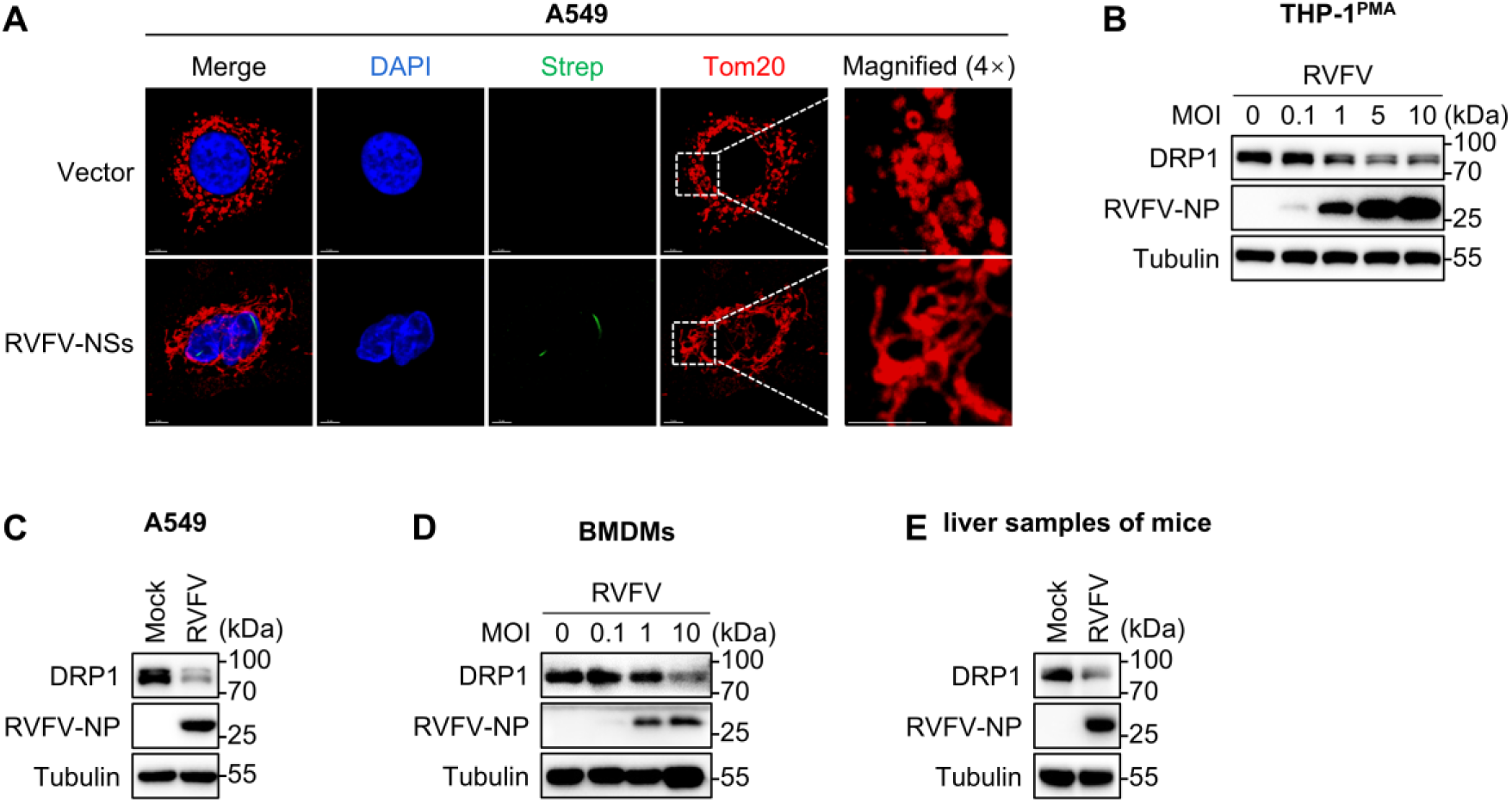
RVFV infection leads to mitochondrial elongation through down-regulating DRP1. **Related to Figure 1.** (A) The effects of exogenously expressed RVFV-NSs protein on mitochondrial morphology in A549 cells was observed by confocal microscopy. A549 cells were transfected with empty vector or vector expressing strep-tagged RVFV-NSs for 48 hours. Mitochondria were labeled with anti-Tom20 antibody (red), RVFV-NSs were labeled with anti-strep antibody (green) and Nuclei were stained with DAPI (blue). Scale bar, 5 μm. (B) Immunoblot analysis of DRP1 protein level in THP-1^PMA^ cells infected with RVFV with indicated MOI. (C) Immunoblot analysis of DRP1 protein level in A549 cells infected with RVFV for 24 h. (D) Immunoblot analysis of DRP1 protein level in mouse bone marrow-derived macrophages (BMDMs) infected with RVFV for 18 h. (E) Immunoblot analysis of DRP1 protein level in the liver samples from the mice infected with RVFV at 1000 PFU for 3 days. Data are representative of three experiments with similar results.

**Figure S2.**
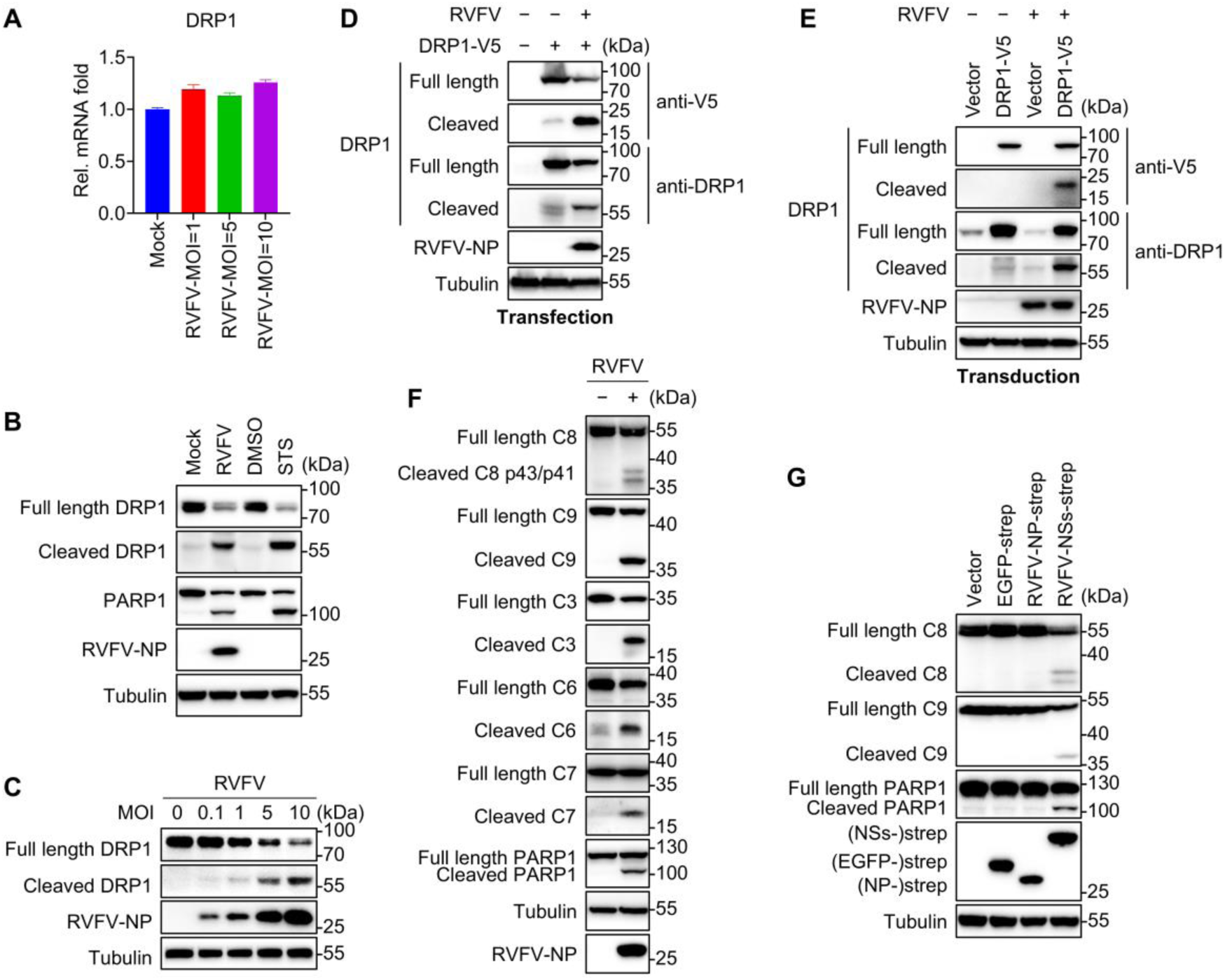
DRP1 is cleaved by activated caspase during RVFV infection. **Related to Figure 2.** (A) qPCR analysis of DRP1 mRNA in THP-1^PMA^ cells infected with RVFV at the indicated MOI for 24 h. (B) Immunoblot analysis of DRP1 cleavage in THP-1^PMA^ cells infected or mock infected with RVFV for 24 h, or treated or untreated with Staurosporine (STS) at 1μM for 12 h. (C) Immunoblot analysis of DRP1 cleavage in THP-1^PMA^ cells infected with RVFV at the indicated MOI. (D) Immunoblot analysis of the DRP1 cleavage in Huh7 cells transfected with empty vector or vector expressing V5-tagged DRP1 (DRP1-V5) for 24 h, followed by RVFV infection or mock infection for 24 h. The DRP1-V5 was detected by anti-V5 antibody or anti-DRP1 antibody, respectively. (E) Immunoblot analysis of the DRP1 cleavage in THP-1^PMA^ cells, transduced with empty vector or vector expressing V5-tagged DRP1 (DRP1-V5), infected or mock infected with RVFV for 24 h. The DRP1-V5 was detected by anti-V5 antibody or anti-DRP1 antibody as indicated. (F) Immunoblot analysis of caspases activation in THP-1^PMA^ cells infected or mock infected with RVFV^WT^ for 24 h. (G) Immunoblot analysis of caspases activation in Huh7 cells transfected with empty vector or vector expressing the indicated proteins for 48 h. Data are representative of three experiments with similar results.

**Figure S3.**
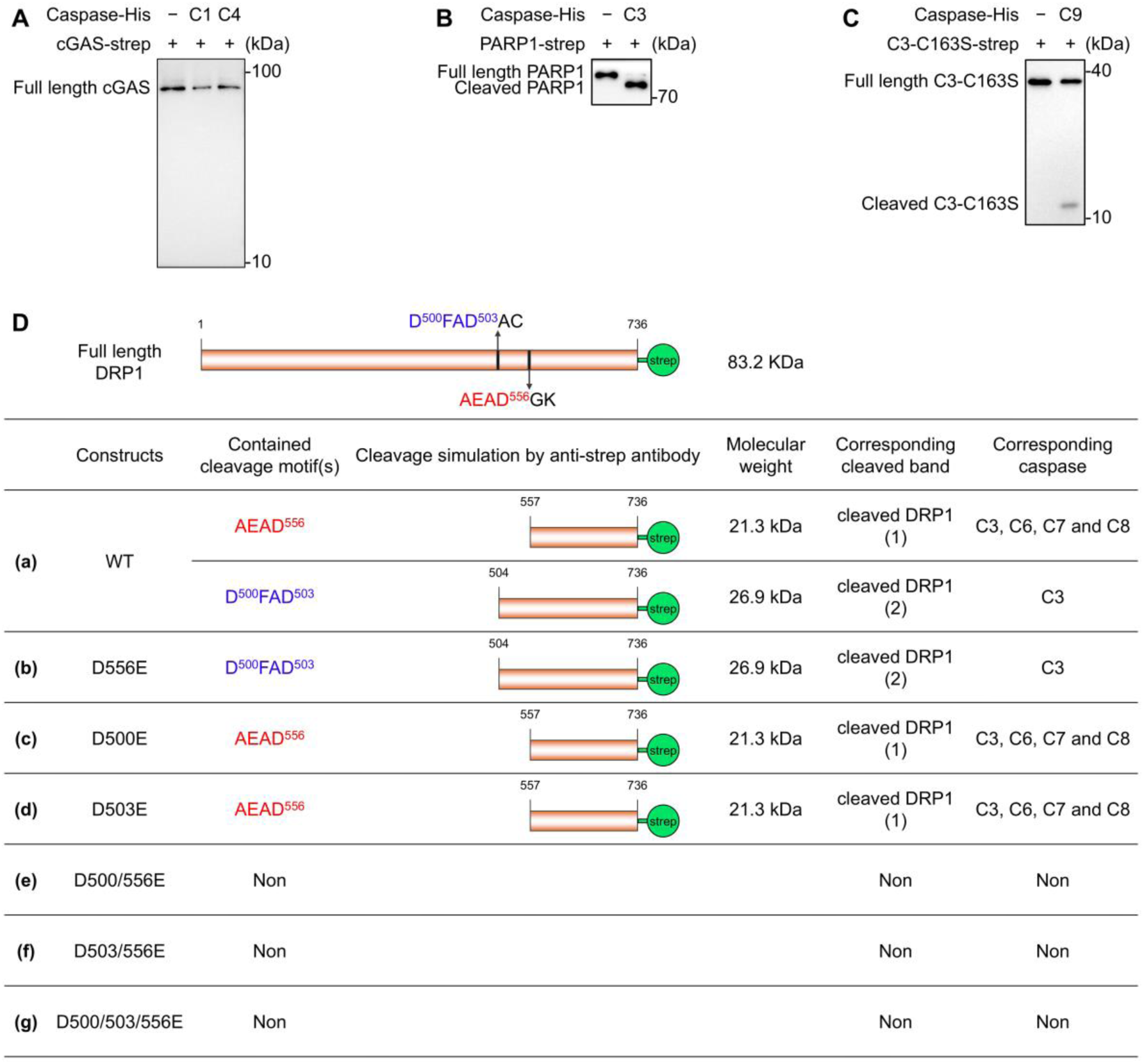
*In vitro* caspase activity validation and schematic of the caspase-mediated DRP1 cleavage. **Related to Figure 3.** (A-C) *In vitro* caspase activity assays. Recombinant caspase activity was confirmed using their respective positive control substrates: (A) cGAS for caspase-1 and -4, (B) PARP1 for caspase-3, and (C) a catalytically inactive mutant of procaspase-3 (C3-C163S) for caspase-9. (D) Schematic model of caspase-mediated cleavage of the C-terminal strep-tagged DRP1, as detected by anti-strep antibody. Related to Figure 4F. (a) DRP1-WT can be cleaved by caspase-3, -6, -7 and -8 at AEAD^556^ motif to generate cleaved DRP1 (1) or cleaved by caspase-3 at D^500^FAD^503^ motif to generate cleaved DRP1 (2). (b) DRP1-D556E mutant can be cleaved by caspase-3 at D^500^FAD^503^ motif to generate cleaved DRP1 (2). (c-d) DRP1-D500E (c) or DRP1-D503 (d) mutant can be cleaved by caspase-3, -6, -7 and -8 at AEAD^556^ motif to generate cleaved DRP1 (1). (e-g) DRP1-D500/556E (e), DRP1-D503/556E (f) or DRP1-D500/503/556E (g) mutant cannot be cleaved by caspase-3, -6, -7 and -8 at AEAD^556^ or D^500^FAD^503^ motif to generate cleaved DRP1 (1) or cleaved DRP1 (2).

**Figure S4.**
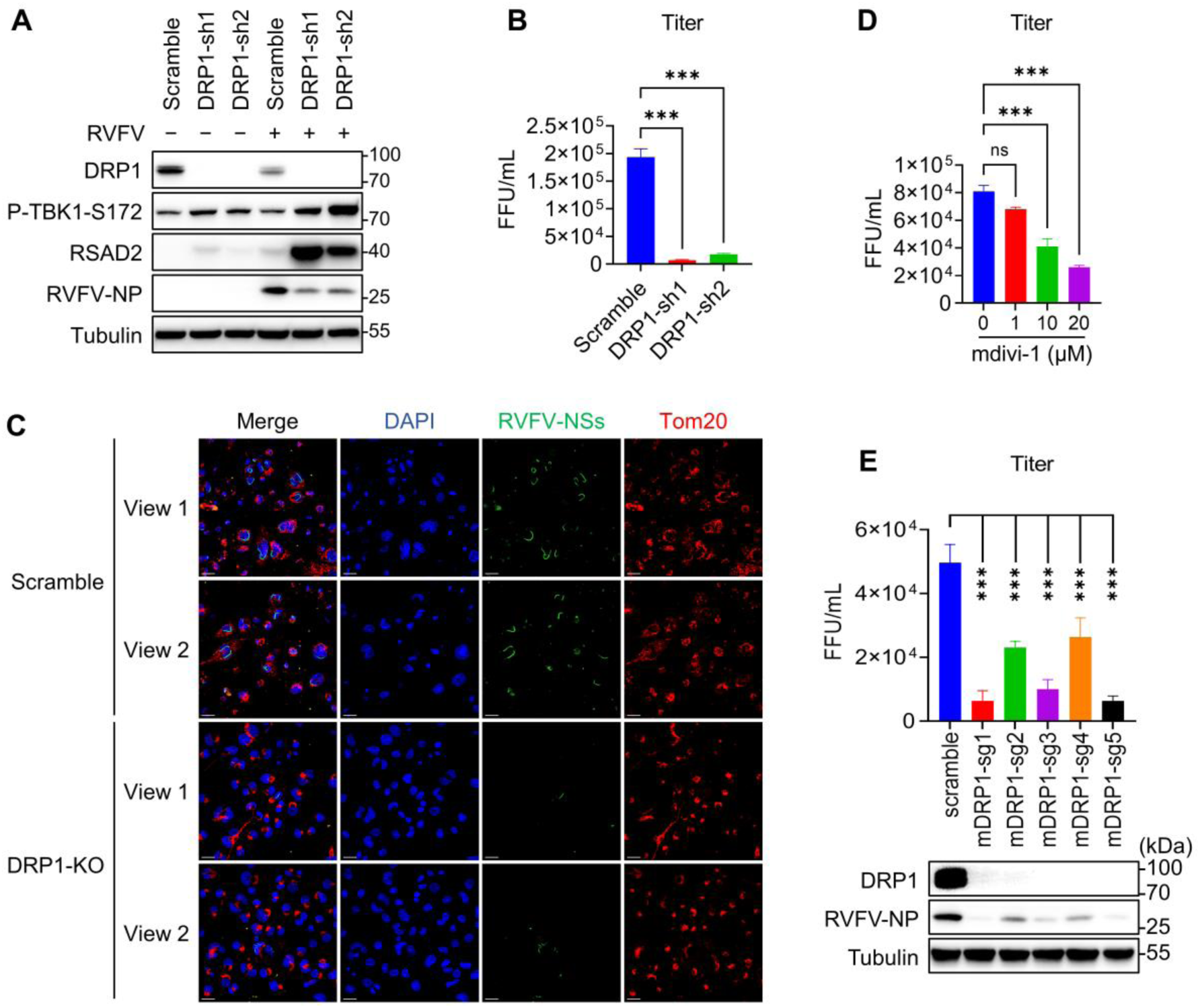
DRP1 deficiency promotes antiviral innate immunity. **Related to Figure 6.** (A) Immunoblot analysis of the indicated proteins in scramble or DRP1 knockdown THP-1^PMA^ cells infected or mock infected with RVFV for 24 h. (B) Focus forming assay to determine the titer from RVFV infected scramble or DRP1 knockdown THP-1^PMA^ cells infected for 24 h. (C) Immunofluorescence analysis of RVFV-NSs expression in scramble or DRP1 knockout THP-1^PMA^ cells infected with RVFV for 24 h. Mitochondria were labeled with anti-Tom20 antibody (red), RVFV-NSs was labeled with anti-RVFV-NSs antibody (green) and Nuclei were stained with DAPI (blue). Scale bars, 20 μm. (D) Focus forming assay to determine the titer from THP-1^PMA^ cells infected with RVFV for 2 h, followed by mdivi-1 at the indicted concentrations for 20 h. (E) Focus forming assay to determine the titer from RVFV infected scramble or DRP1 knockout MEF cells. Cells were infected for 24 h, and cell lysate was harvested for immunoblot analysis. \*\**P* < 0.01, \*\*\**P* < 0.001; ns, not significant (One-way ANOVA test). Data are representative of three experiments with similar results.

**Figure S5.**
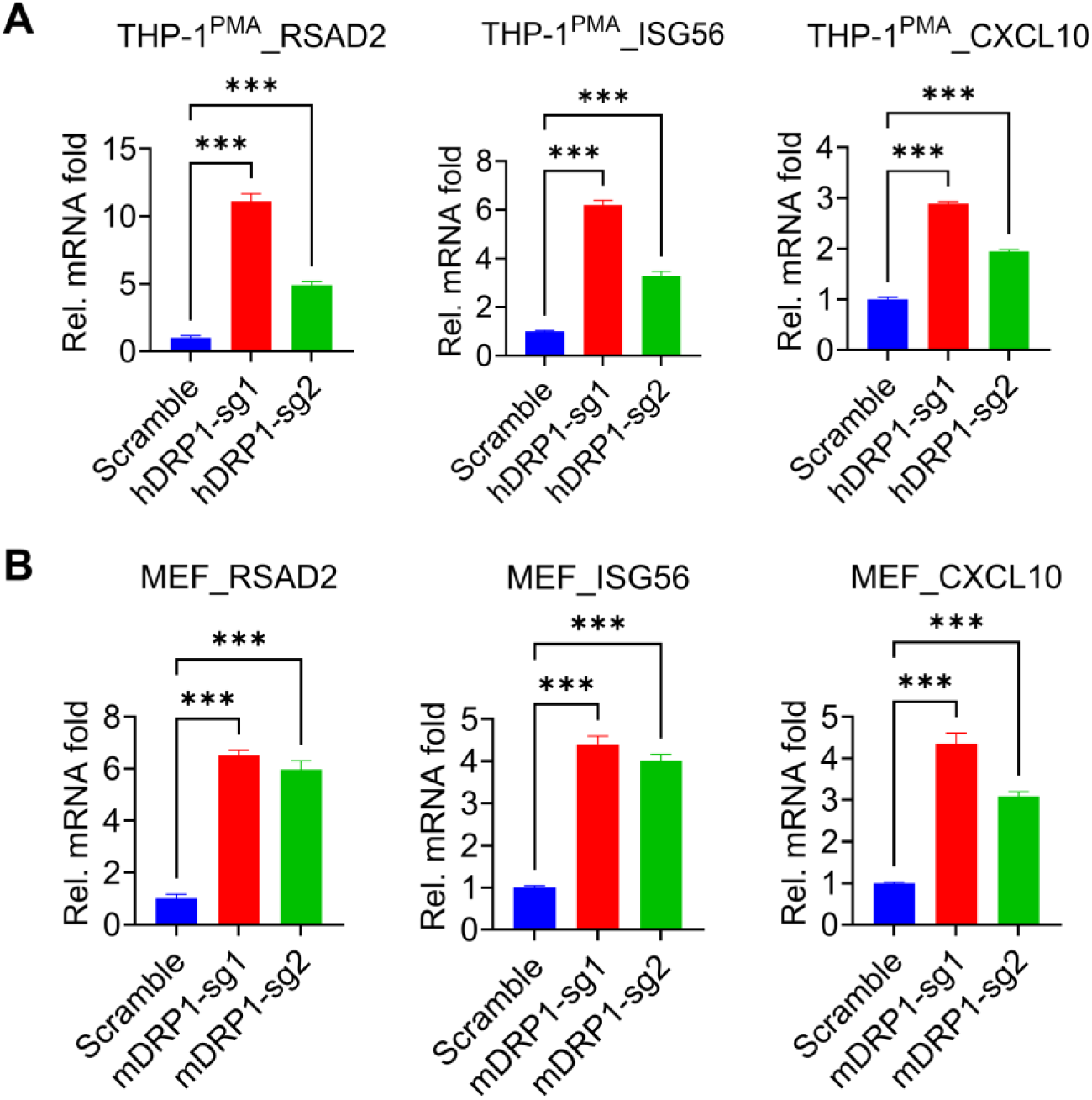
DRP1 deficiency promotes immune responses. **Related to Figure 7.** (A-B) qPCR analysis of the indicted genes in scramble or DRP1 knockout THP-1^PMA^ cells (A) or MEF cells (B). \*\**P* < 0.01, \*\*\**P* < 0.001 (One-way ANOVA test). Data are representative of three experiments with similar results.

**Table S1.**
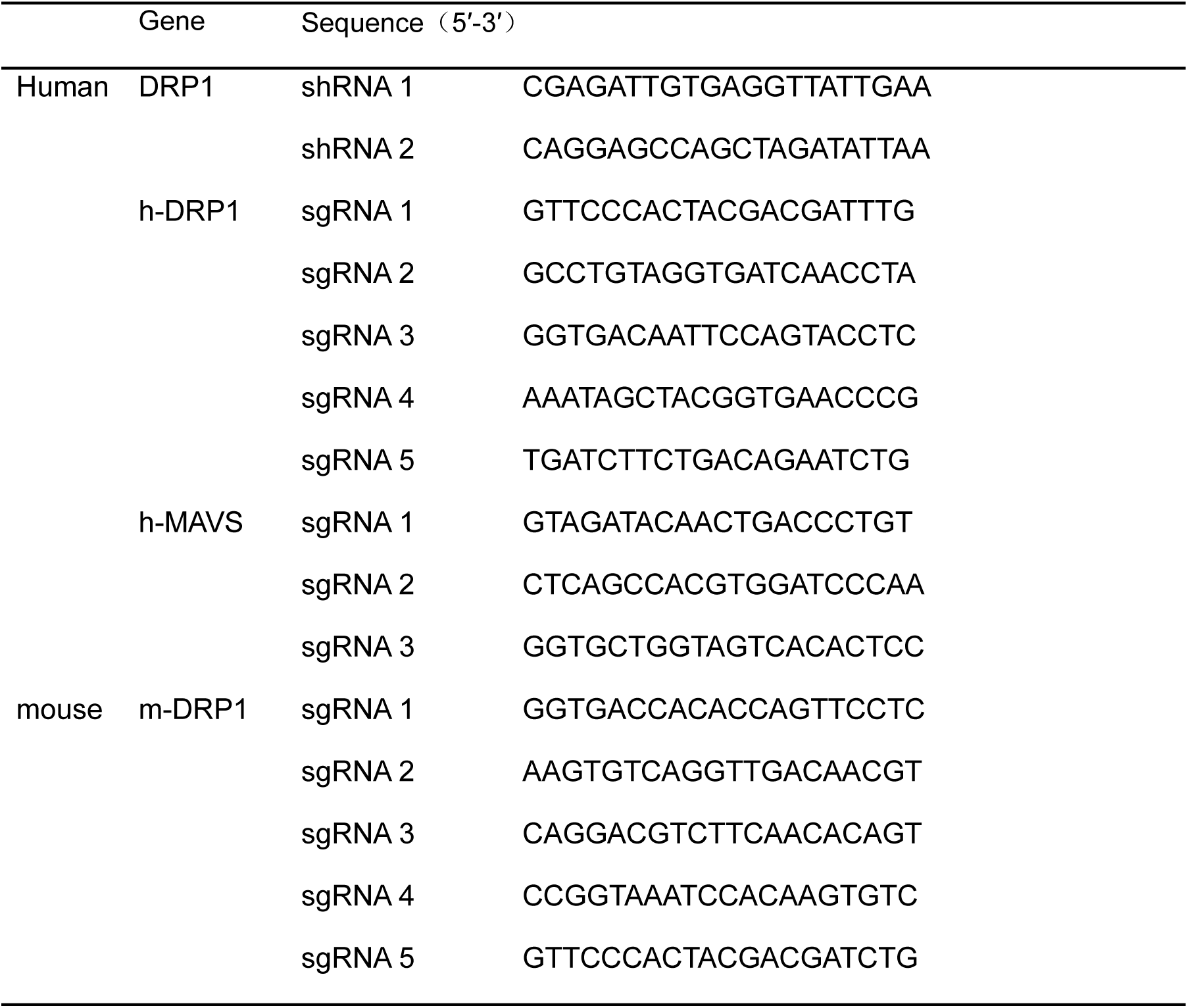
The sequences of corresponding shRNA or sgRNA sequence.

**Table S2.**
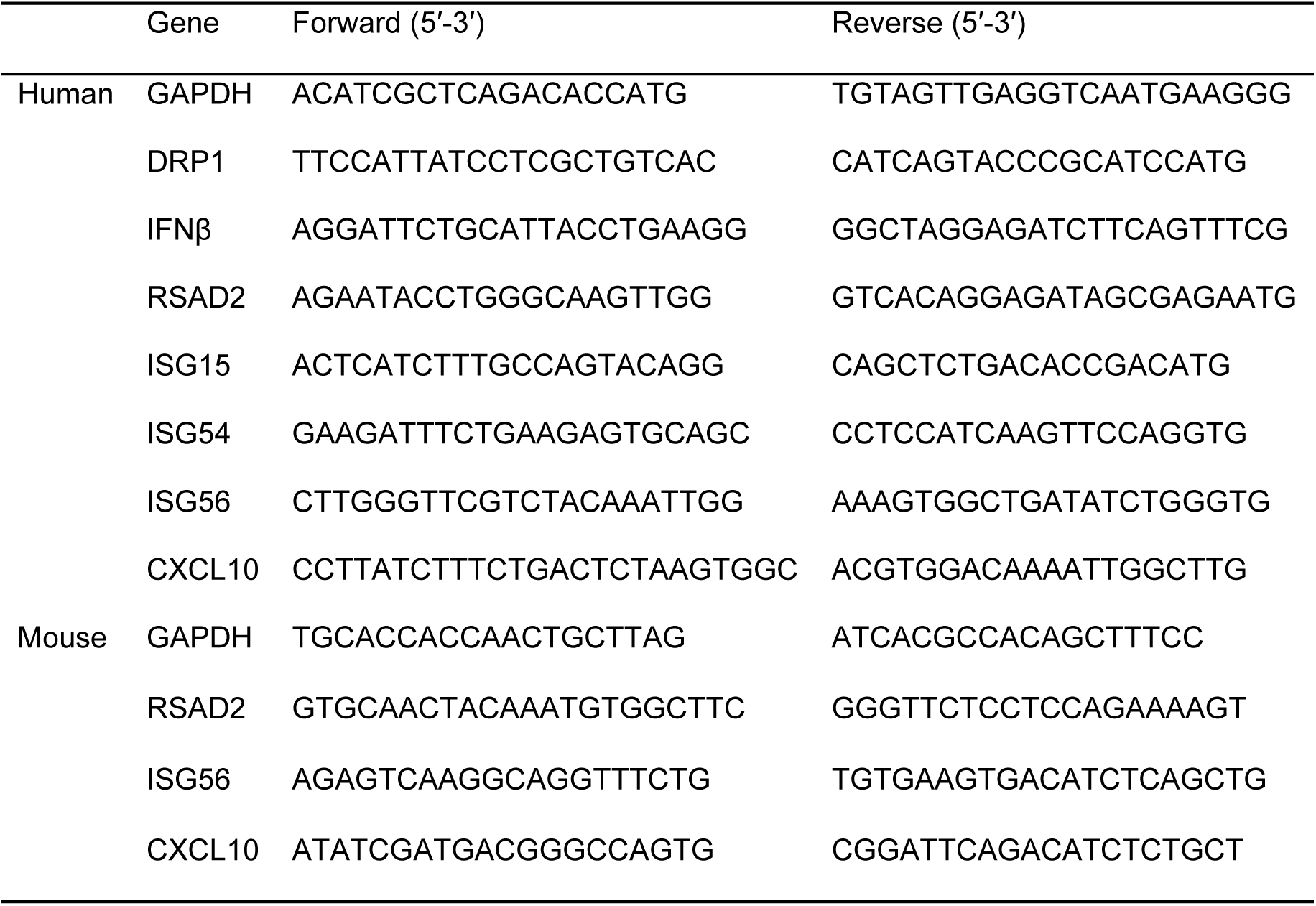
The sequences of gene-specific primers.

